# No Evidence of Direct Activation of Human Neutrophil Responses by Multivalent Prefusion Trimeric SARS-CoV-2 Spike Protein *ex vivo*

**DOI:** 10.1101/2025.05.08.652856

**Authors:** Audray Fortin, Sandrine Huot, Elise Caron, Cynthia Laflamme, Amélie Pagliuzza, Nicolas Chomont, Caroline Gilbert, Baoshan Zhang, Peter D. Kwong, Marc Pouliot, Nathalie Grandvaux

## Abstract

The SARS-CoV-2 Spike (S) protein is essential for viral entry and serves as the primary immunogen in most COVID-19 vaccines. While its role in adaptive immunity is well defined, its potential to contribute directly to innate immune activation and remains incompletely understood. Neutrophils, in particular, are prominent effectors in COVID-19 severity, yet how they respond directly to the S protein presented in a multivalent format is unclear. Here, we investigated whether the S protein can directly activate human neutrophils *ex vivo* using two biologically relevant models: nanoparticles displaying multivalent stabilized prefusion trimeric S glycoprotein, and purified β-propiolactone-inactivated SARS-CoV-2 virions. Neutrophils were exposed to nanoparticles or inactivated virus, either alone or pre-coated with monoclonal or polyclonal anti-S antibodies. Nanoparticles displaying Respiratory Syncytial Virus (RSV) Fusion (F) protein and purified β-propiolactone-inactivated RSV served as comparators. Across all models and conditions tested, the S protein did not induce significant neutrophil responses. No consistent effects were observed on cell viability, surface marker expression, reactive oxygen species production, neutrophil extracellular trap formation, cytokine release, or inflammatory gene expression—even in the presence of anti-S antibodies mimicking immune complexes. Results with F-nanoparticles and inactivated RSV were similarly modest. These findings indicate that the trimeric prefusion S protein, whether displayed multivalently on nanoparticles or in the context of inactivated viral particles, is insufficient to trigger robust neutrophil activation. This work provides insight into the innate immune profile of the S protein and suggests that its use in vaccine platforms is unlikely to directly provoke neutrophil-mediated inflammatory responses.

## Introduction

In December 2019, the emergence of severe acute respiratory syndrome coronavirus 2 (SARS-CoV-2) led to a global pandemic, profoundly impacting global health and economies. Despite extensive research, the factors driving the wide spectrum of clinical manifestations of SARS-CoV-2 infection—from asymptomatic cases to severe disease with multi-organ failure and cardiovascular and neurological complications—remain incompletely understood [1]. It is well established, however, that severe COVID-19 is often associated with an excessive inflammatory response, commonly referred to as a cytokine storm, which contributes to tissue damage and systemic complications [2, 3]. Beyond the acute phase, a significant proportion of individuals develop post-acute sequelae of COVID-19 (PASC), also known as Long COVID, characterized by persistent symptoms that can last for months or even years [4]. A growing body of evidence suggests that Long COVID symptoms are driven by the existence of a persisting SARS-CoV-2 reservoir in a subset of affected patients [5, 6].

Neutrophils play a pivotal role in the innate immune response, acting as first-line defenders against infections through mechanisms such as migration to infection sites, reactive oxygen species (ROS) production, neutrophil extracellular trap (NET) formation, and cytokine release [7]. However, excessive, or dysregulated neutrophil activation can lead to oxidative stress, tissue damage, and systemic inflammation. In the context of COVID-19, neutrophils have been strongly implicated in disease pathogenesis [8–10]. Severe COVID-19 is associated with increased neutrophil counts in peripheral blood, distinct neutrophil activation patterns predictive of critical illness, and neutrophil-driven immunopathology in affected organs [11–13]. Furthermore, elevated levels of NETs have been reported in COVID-19 patients, correlating with disease severity. SARS-CoV-2 infection induces NET formation in vitro, and post-mortem analyses reveal NET accumulation in the lungs of fatal COVID-19 cases, suggesting a role in acute respiratory distress syndrome (ARDS) and thromboinflammation [14–16]. Despite these insights, the precise molecular triggers underlying neutrophil activation in COVID-19 remain incompletely defined.

Among the viral components implicated in disease pathogenesis, the SARS-CoV-2 Spike (S) glycoprotein is of particular significance. The S protein facilitates viral entry by binding to the angiotensin-converting enzyme 2 (ACE2) receptor on host cells [17]. In its functional prefusion state, S is a trimeric protein composed of S1 and S2 subunits, with cleavage at the furin site separating the two domains. S1 mediates receptor engagement, while S2 undergoes conformational changes to drive membrane fusion [18–20]. As the principal target of neutralizing antibodies, the SARS-CoV-2 S protein has been central to vaccine development, pursued through various strategies, including mRNA-based (BNT162b2, Pfizer-BioNTech; mRNA-1273, Moderna), adenoviral vector-based (AZD1222/ChAdOx1 nCoV-19, AstraZeneca; Ad26.COV2.S, Janssen; Sputnik V, Gamaleya), and protein subunit vaccines (NVX-CoV2373, Novavax). With the exception of AZD1222/ChAdOx1 and Sputnik V, these vaccines are designed to elicit immune responses against a stabilized prefusion form of S, incorporating stabilizing mutations to prevent structural conversion and thereby optimize antigenicity and immunogenicity [21]. Inactivated virus-based vaccines (CoronaVac, Sinovac; BBIBP-CorV, Sinopharm; BBV152, Bharat Biotech) also aim to present the native prefusion S as an immunogen; however, post-fusion conformations may arise during purification [21]. Despite the occurrence of rare adverse effects, including vaccine-induced immune thrombotic thrombocytopenia (VITT) associated with AZD1222/ChAdOx1 nCoV-19 and Ad26.COV2.S adenoviral vector vaccines, vaccines have demonstrated a strong benefit-risk profile, effectively preventing severe disease, reducing hospitalizations, and mitigating the global impact of COVID-19 [22, 23].

Beyond its role in viral entry, emerging evidence suggests that the S protein itself may contribute to COVID-19 immunopathology. Multiple pieces of evidence support that S can induce microvascular dysfunction, disrupt the blood-brain barrier, and trigger inflammatory responses in microglia and other immune cells [24–26]. Furthermore, consistent with the presence of a SARS-CoV-2 reservoir, the S protein has been detected in tissues and circulation long after acute infection, suggesting a potential role in sustained inflammation and post-acute sequelae [27–29]. While rare, adverse effects of COVID-19 vaccines such as myocarditis and thrombotic events share similarities with severe COVID-19 complications. This has led to the speculation that the S protein detected in the plasma of vaccine recipients, particularly those with myocarditis, may contribute to inflammation-related adverse reactions [30–33]. Finally, while there is little clinical evidence of antibody-dependent enhancement of disease, SARS-CoV-2 S-specific IgG have been linked to excessive inflammatory responses in COVID-19 [34].

Despite previous research on neutrophils and the SARS-CoV-2 S protein, the functional interaction of the trimeric prefusion glycoprotein in a multivalent format remains poorly understood. While most available studies have used soluble recombinant S protein to assess neutrophil activation, here we investigated whether the S protein can directly activate human neutrophils *ex vivo* using two biologically relevant models: a stabilized prefusion trimeric S protein displayed multivalently on nanoparticles, and β-propiolactone-inactivated SARS-CoV-2 virions. Strikingly, our findings reveal little to no direct neutrophil activation in response to the S protein, challenging the notion that S is sufficient to trigger robust neutrophil activation.

## Materials and Methods

### Materials

Dextran-500, Phorbol 12-myristate 13-acetate (PMA), Luminol, β-propiolactone (BPL) and PEG-6000 were purchased from Sigma-Aldrich (Oakville, ON, Canada). Lymphocyte separation medium was from Wisent (St-Bruno, QC, Canada). Lyophilized human immunoglobulins G (IgGs), purified from human plasma or serum by fractionation (purity >97% determined by SDS-PAGE), were from Innovative Research, Inc (Novi, MI, USA). Fc Block™ was from BD Biosciences (San Jose, CA, USA). SYTOX™ Green and CellTrace™ Violet were from Thermo Fisher Scientific (Burlington, ON, Canada).

### Nanoparticles displaying trimeric prefusion SARS-CoV-2 S or RSV-F protein

Self-assembling nanoparticles with a lumazine synthase (LuS)-based scaffold with multivalent display of viral glycoprotein used in this study were produced and validated as we previously described in [35]. The LuS-N71-SpyLinked-RSV F nanoparticles (F-nanoparticles; M.W. 2584.2 kDa) display prefusion-stabilized trimeric respiratory syncytial virus (RSV) fusion (F) glycoprotein (DS2-preF stabilized RSV F). Similarly, the LuS-N71-SpyLinked-CoV-2 S nanoparticles (S-nanoparticles; M.W. 6260.1 kDa) present the prefusion-stabilized trimeric version (GSAS and PP mutations and the T4 phage fibritin trimerization domain) of the SARS-CoV-2 S protein.

#### SARS-CoV-2 amplification and purification

All experiments involving SARS-CoV-2 culture were conducted in a certified containment level-3 facility following standard operating procedures approved by the Biosafety Committee at CRCHUM, Montreal, Canada. The SARS-CoV-2/SB2 isolate [36] was obtained from Dr. Samira Mubareka (Sunnybrook Research Institute, Toronto, Canada) and propagated in VERO E6 cells (ATCC) as described in [37]. Viral stocks were sequenced and tested for mycoplasma contamination (Invivogen). SARS-CoV-2 was amplified in VERO E6 cells in DMEM (GIBCO) supplemented with 1% L-glutamine and 2% Fetaclone III (Hyclone) at a multiplicity of infection (MOI) of 0.02. At 96 h post-infection, the supernatant was harvested and clarified by centrifugation at 4000g for 15 min to remove cell debris. Viral inactivation was achieved by treatment with 0.05% (v/v) β-propiolactone (BPL) for 16 h at 4°C, followed by hydrolysis at 37°C for 2 h. Inactivation was confirmed by the absence of viral replication in VERO E6 cells, quantified using the median tissue culture infectious dose (TCID50) method [37]. The clarified inactivated supernatant was precipitated by gentle stirring on ice for 90 min with 10% PEG-6000 in the presence of 100 mM MgSO₄. The pellet was dissolved in cold NTE buffer (150 mM NaCl, 50 mM Tris-HCl pH 7.5, 1 mM EDTA) and further purified using a discontinuous sucrose gradient (10%, 20%, and 30% sucrose in NTE). Ultracentrifugation was performed at 41,000 rpm (211,400 g) for 60 min at 4°C using a TH-641 Swinging Bucket Rotor (Thermo Scientific™). The purified virus pellet was gently resuspended in a minimal volume of cold NTE buffer to obtain a high-concentration virus stock, which was aliquoted and stored at -80°C. Viral fractions were resolved by SDS-PAGE and immunoblotted as previously described [38], using anti-SARS-CoV-2 S (BEI Resources, NR-52947) and anti-SARS-CoV-2 N (Novus, NB100-56576) antibodies. Virion quantification was performed via RT-qPCR. RNA extraction from virus preparations was carried out using the QIAamp Viral RNA Mini Kit (Qiagen), and the *SARS-CoV-2 N* gene was quantified by qPCR on a QuantStudio 5 instrument using TaqPath 1-Step Multiplex Master Mix No ROX (Applied Biosystems). The following primers and probe were used: N_Forward: 5’-CGTACTGCCACTAAAGCATACA-3’; N_Reverse: 5’-GCGGCCAATGTTTGTAATCAG-3’; N_P: 5’-AGACGTGGTCCAGAACAAACCCAA-3’. A standard curve was generated using *in vitro* transcribed RNA, and the quantity of SARS-CoV-2 particles per mL was extrapolated based on *N* gene quantification.

#### RSV A2 amplification and purification

All experiments involving RSV culture were conducted in a certified containment level-2 facility following standard operating procedures approved by the Biosafety Committee at CRCHUM, Montreal, Canada. The initial stock of the RSV A2 strain was obtained from Advanced Biotechnologies Inc. Virus amplification was performed in HEp-2 cells (ATCC) cultured in DMEM (GIBCO) supplemented with 1% L-glutamine (GIBCO) and 2% Fetaclone III (Hyclone) at a multiplicity of infection (MOI) of 0.1, until 50% cytopathic effect was observed. Cells and culture medium were harvested by scraping, followed by centrifugation at 3200 g for 20 min at 4°C. The resulting pellet underwent three freeze-thaw cycles before a final centrifugation step. Supernatants were pooled and inactivated by treatment with 0.05% (v/v) β-propiolactone (BPL) for 16 h at 4°C, followed by hydrolysis at 37°C for 2 h. Inactivated RSV was then precipitated with 10% PEG-6000 as described in [39] and resuspended in 20% sucrose in NTE buffer using a Dounce homogenizer. The virus suspension was layered onto a 30% sucrose cushion in NTE and ultracentrifuged at 25,000 rpm (106,000 g) at 4°C for 1 h using an SW41 Ti rotor (Beckman Coulter). The resulting pellet was resuspended in 5% sucrose in NTE using a Dounce homogenizer, aliquoted, and stored at -80°C until use. Virus inactivation was confirmed by methylcellulose plaque assays, as previously described [40]. Purified RSV was resolved by SDS-PAGE and immunoblotted using anti-RSV antibodies (Chemicon, AB1128). Virion quantification (particles per mL) was determined based on RSV A2 *N* gene quantification by RT-qPCR. RNA extraction and qPCR conditions were as described for SARS-CoV-2, except that amplification of the *N* gene was performed on a Rotor-Gene 3000 Real-Time Thermal Cycler (Corbett Research) using the FastStart SYBR Green Kit (Roche) with the following primers: N_Forward: 5’-AGATCAACTTCTGTCATCCAGCAA-3’; N_Reverse: 5’-TTCTGCACATCATAATTAGGAGTATCAAT-3’.

#### Hydrodynamic size measurement

The size and homogeneity of nanoparticles and inactivated-viral particles size was analyzed by dynamic light scattering (DLS) with a Zetasizer Nano ZS (Malvern Instruments) as previously described in [41]. Hydrodynamic diameter measurements were done in duplicate at room temperature.

#### Coating of nanoparticles and viruses with antibodies

Where indicated, S- and F-nanoparticles were pre-coated with specific antibodies against the S- or F-proteins, respectively, for 30 min at 37°C. For S-nanoparticles, a neutralizing monoclonal anti-S antibody (Clone CV30; Absolute Antibody, Boston, USA) was used. In subset experiments, purified SARS-CoV-2 virions were coated either with CV30 alone (αS) or with a combination of five monoclonal antibodies (αS Mix), each targeting a distinct epitope on the SARS-CoV-2 S protein RBD domain (Leinco Technologies Inc., St. Louis, MO, USA). At least one of these antibodies possesses neutralizing activity. For F-nanoparticles, Palivizumab (Synagis, AbbVie), a recombinant humanized monoclonal IgG1 anti-F antibody was used [42]. The details of the anti-S and anti-F antibodies used in this study are provided in **Table 1**.

**Table 1.**
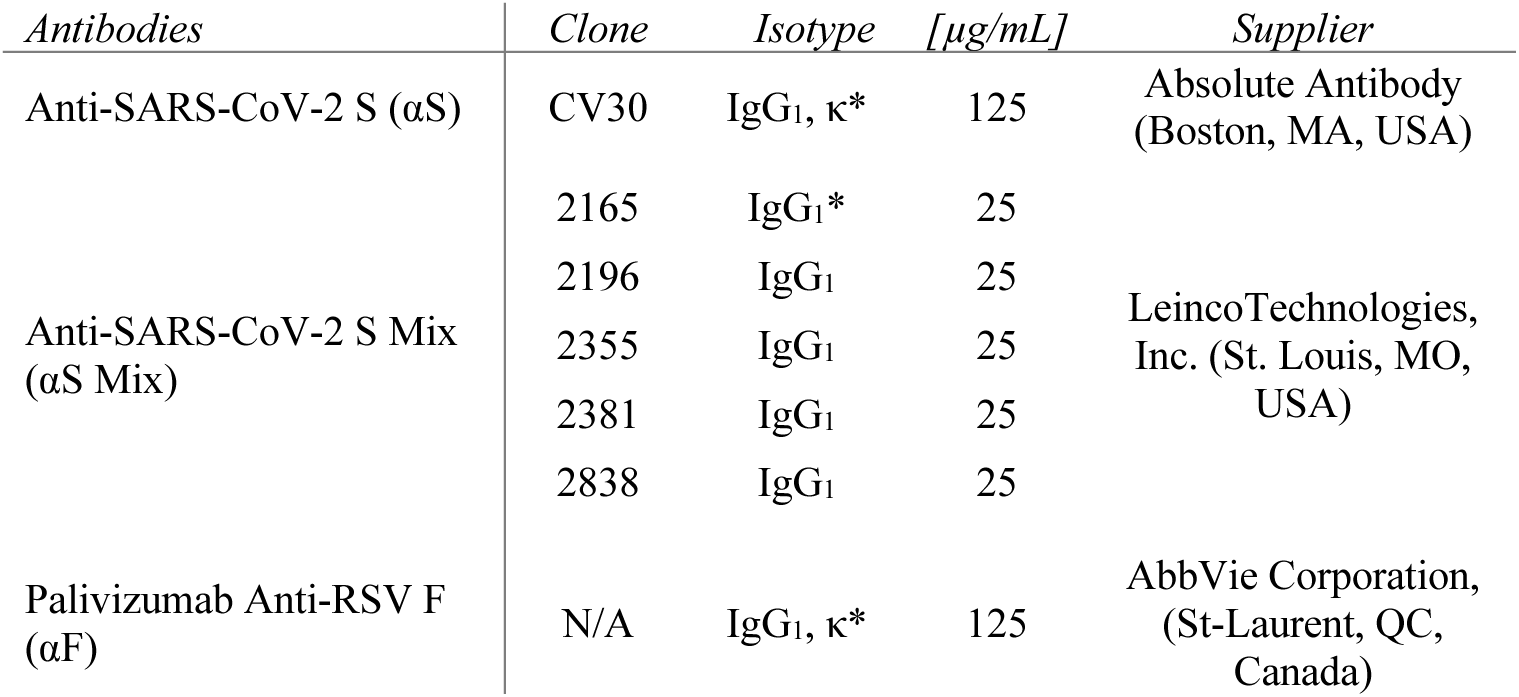
Monoclonal antibodies used in this study. All anti-S antibodies target the RBD. *\*Neutralizing antibodies*.

#### Human neutrophils isolation

Written informed consent was obtained from all donors. Data collection and analysis were performed anonymously. Neutrophils were isolated as previously described, with modifications [43]. Briefly, venous blood from healthy volunteers was collected in isocitrate anticoagulant solution and centrifuged at 250 g for 10 min to separate the platelet-rich plasma, which was discarded. Leukocytes were obtained following erythrocyte sedimentation in 2% Dextran-500. Neutrophils were then separated from other leukocytes by centrifugation through a 10 mL layer of lymphocyte separation medium. Residual erythrocytes were eliminated by a 20-second hypotonic lysis step. The final granulocyte preparation consisted of >95% neutrophils and <5% eosinophils, with monocyte contamination below 0.2%, as determined by esterase staining. Cell viability exceeded 98%, as assessed by trypan blue dye exclusion. All isolation steps were performed under sterile conditions at room temperature.

#### Preparation of heat-aggregated (HA) IgGs

HA-IgGs were prepared as previously described [44], with modifications. Briefly, soluble aggregates were freshly generated each day by resuspending IgGs in HBSS at a concentration of 25 mg/mL, followed by heating at 63°C for 75 min.

#### Neutrophil incubation with nanoparticles or inactivated viruses

Neutrophils were resuspended in Hank’s Balanced Salt Solution (HBSS) supplemented with 10 mM HEPES (pH 7.4) and 1.6 mM Ca²⁺, but without Mg²⁺. The nanoparticle-to-neutrophil and virus particle-to-neutrophil ratios are provided in each figure.

#### Neutrophil viability assessment

Neutrophil viability was evaluated using the LIVE/DEAD™ Fixable Dead Cell Stain Kit (Thermo Fisher Scientific, Burlington, ON, Canada). Briefly, cell pellets were resuspended in 1 mL of HBSS containing 1 μL of LIVE/DEAD stain and incubated for 30 min. Samples were then centrifuged, resuspended in 400 μL of 1% paraformaldehyde, and analyzed by flow cytometry.

#### Analysis of neutrophil surface marker expression

Following treatment, cell suspensions were centrifuged and resuspended in HBSS containing human Fc Block™ (BD Biosciences). Cells were incubated with the appropriate mix of fluorophore-conjugated antibodies for 30 min at 4°C in the dark. After incubation, cells were centrifuged, resuspended in 400 μL of 1% paraformaldehyde, and analyzed by flow cytometry using a FACS Canto II instrument with FACSDiva software (version 6.1.3, BD Biosciences). A detailed list of labeled mouse monoclonal antibodies targeting the surface markers CD15, CD46, CD55, CD59, CD64, CD93, CD32, CD16, CD62L, CD184, CD63, CD66b, and CD11b (all from BD Biosciences, San Jose, CA, USA) is provided in **Table 2**.

**Table 2.**
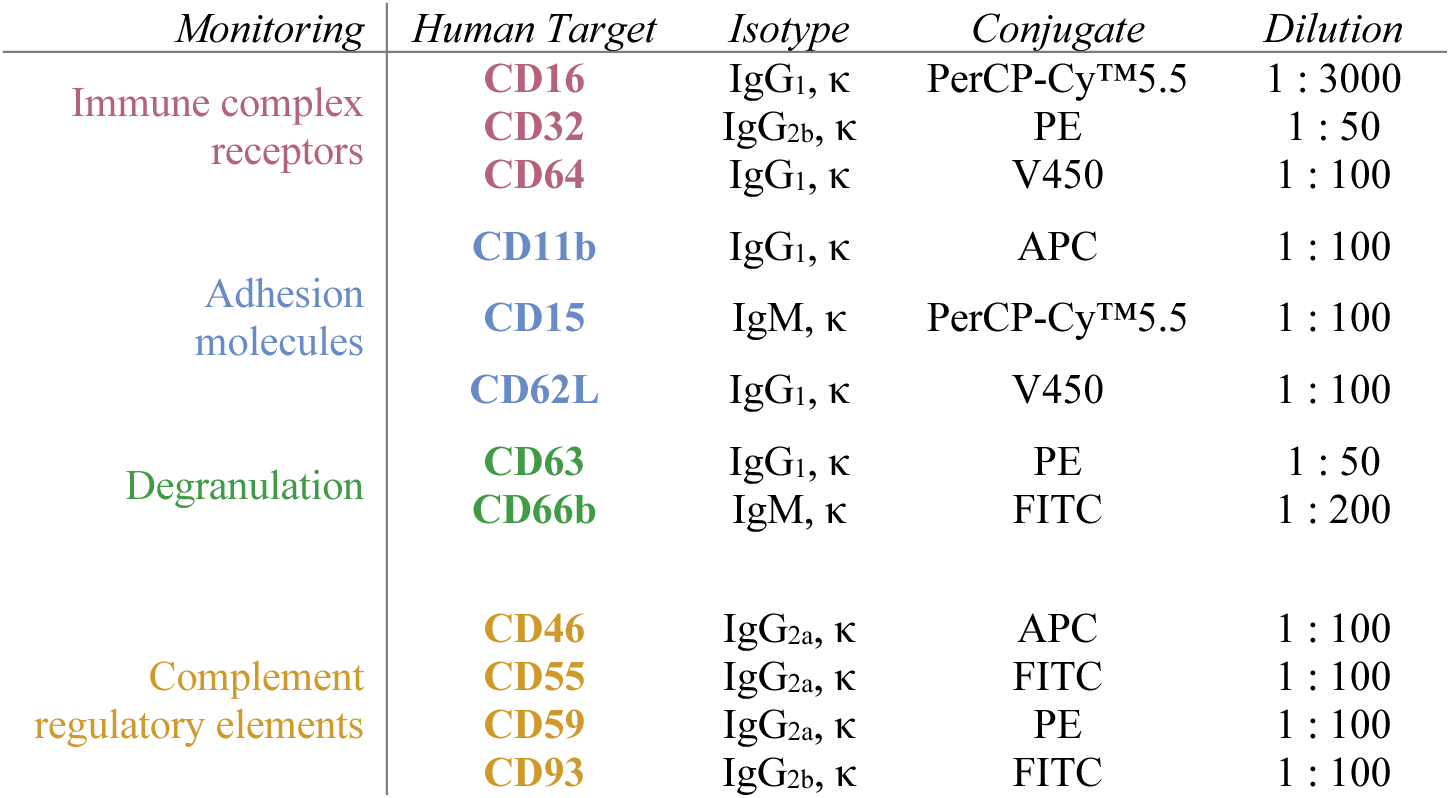
List and description of mouse monoclonal antibodies used in flow cytometry to monitor surface markers.

#### Measurement of ROS production by neutrophils

ROS production was measured as previously described [45], with modifications. Briefly, neutrophils were resuspended at 1 × 10⁶ cells/mL in HBSS supplemented with 10% heat-inactivated fetal bovine serum (FBS) and 10 μM luminol. Neutrophils (200 μL) were seeded into 96-well microplates, treated as indicated, and stimulated with or without 1 mg/mL HA-IgGs. Suspensions were incubated at 37°C in an Infinite M1000 PRO microplate reader (Tecan, Morrisville, NC, USA) using i-control 2.0 software. Luminescence intensity was recorded every 5 min.

#### Measurement of Neutrophil Extracellular Trap (NET)

NET production was assessed as previously described [46], with modifications. Briefly, neutrophils were resuspended at 2.5 × 10^5^ cells/mL in HBSS, aliquoted into 96-well white plates (200 μL/well) and allowed to settle for 30 min at 37°C. Cells were then treated as indicated and incubated at 37°C with 5% CO₂ for 4 h. SYTOX Green (5 μM final concentration), a cell-impermeable nucleic acid stain, was added, and NET production was quantified by measuring fluorescence at excitation/emission wavelengths of 504/523 nm.

#### Neutrophil RNA Isolation and RT-qPCR Analysis

Total RNA was isolated from neutrophils using Trizol™ Reagent (Thermo Fisher Scientific) following the manufacturer’s protocol, with modifications. Briefly, 25 × 10⁶ neutrophils were homogenized in 1 mL of Trizol, and 200 μL of chloroform was added. After mixing, samples were centrifuged at 12,000 g for 15 min at 4°C. The upper aqueous phase was transferred to a tube containing an equal volume of isopropanol, mixed by inversion, and centrifuged at 12,000g for 10 min at 4°C. The resulting RNA pellets were washed with 1 mL of 75% ethanol, followed by centrifugation at 10,000 g for 5 min at 4°C. After discarding the supernatants, pellets were air-dried for 10–15 min before resuspension in DEPC-treated water. RNA concentration was determined using a Qubit® Fluorometer (Thermo Fisher Scientific).

First-strand cDNA synthesis was performed using 1 µg of total RNA with the SuperScript™ II Reverse Transcriptase (Thermo Fisher Scientific), following the manufacturer’s instructions. Real-time PCR was carried out as described in [47] using SensiFAST SYBR® Lo-ROX mix (FroggaBio, Toronto, ON, Canada) in a Rotor-Gene Q system with Q-series software version 2.0.2 (Qiagen Inc., Mississauga, ON, Canada). Each reaction mixture contained 40 ng of cDNA and 500 nM primers in a final volume of 20 µL. Reaction specificity was confirmed using a melt curve procedure (58–99°C, 1°C per 5 seconds) at the end of the amplification protocol, according to the manufacturer’s instructions. Specific primers for each gene of interest were designed as previously described [48] and are listed in **Table 3**.

**Table 3.**
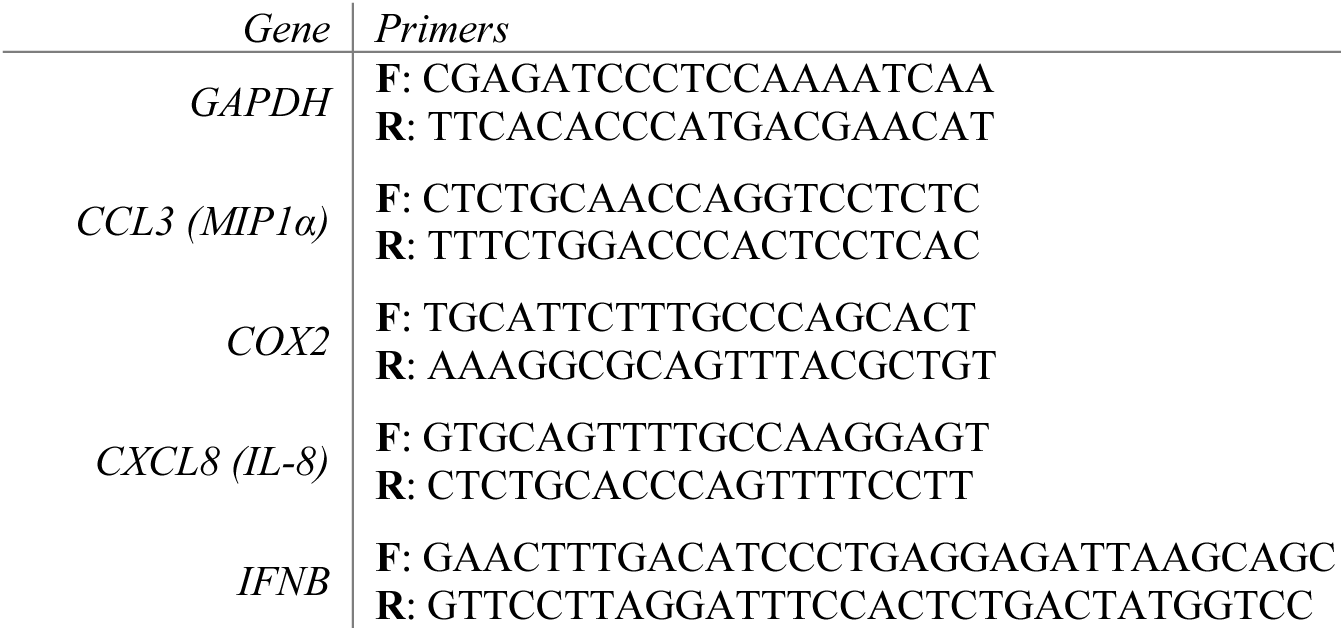
Sequence of PCR primers used for neutrophils mRNA expression analyses. F: forward, R: reverse.

#### Quantification of cytokine and chemokine release from neutrophils

Cell-free supernatants were stored at -20°C until analysis of cytokine and chemokine content using a multiplexed bead-based immunoassay (BD™ Cytometric Bead Array), following the manufacturer’s protocol. Measurements were acquired on a FACS Canto II flow cytometer and analyzed using FCAP Array software (version 3.0, BD Biosciences). The levels of IL-1α, IL-1β, IL-6, CXCL8 (IL-8), TNF, IFNα, CCL3 (MIP-1α), and CCL2 (MCP-1) were quantified.

#### Statistical analysis

Statistical analysis was performed using GraphPad PRISM version 9 (GraphPad Software, San Diego, CA, USA). Where applicable, values are expressed as the mean ± standard error of the mean (SEM). Statistical analysis was performed using the Friedman test followed by a Dunn’s multiple comparisons post hoc test, unless stated otherwise. * *P*<0.05; ** *P*<0.01.

## Results

### Experimental study models

To investigate the influence of the SARS-CoV-2 S protein on human neutrophils *ex vivo*, we employed two distinct experimental systems (**Figure 1**). First, we utilized immunogenic nanoparticles displaying the 2P-stabilized prefusion trimeric SARS-CoV-2 S glycoprotein (LuS-N71-SpyLinked-CoV-2 S nanoparticles, hereafter referred to as S-nanoparticles). As a comparator to determine whether the observed effects were specific to SARS-CoV-2 S or a more general feature of respiratory virus glycoproteins exposed on nanoparticles, we used nanoparticles presenting DS2 stabilized prefusion trimeric respiratory syncytial virus (RSV) fusion (F) glycoproteins (LuS-N71-SpyLinked-RSV F nanoparticles, hereafter referred to as F-nanoparticles). The RSV prefusion F protein is a well-characterized respiratory virus antigen that was used in Arexvy, (GlaxoSmithKline) and Abrysvo (Pfizer) vaccines and successfully elicited protective immune responses [49, 50]. The production of S- and F-nanoparticles has been previously described [35]. These nanoparticles present trimeric glycoproteins in a multivalent format and have been shown to induce significantly stronger neutralizing responses compared to recombinant S protein alone [35]. Therefore, the nanoparticle platform provides a valuable tool for studying the impact of the SARS-CoV-2 S protein in the prefusion trimeric state on neutrophil activation. Characterization of the S- and F-nanoparticle preparations by dynamic light scattering (DSL) revealed a single peak at the expected size ([35], **Figure 1A**), confirming their homogeneity and structural integrity at the time of use.

**Figure 1.**
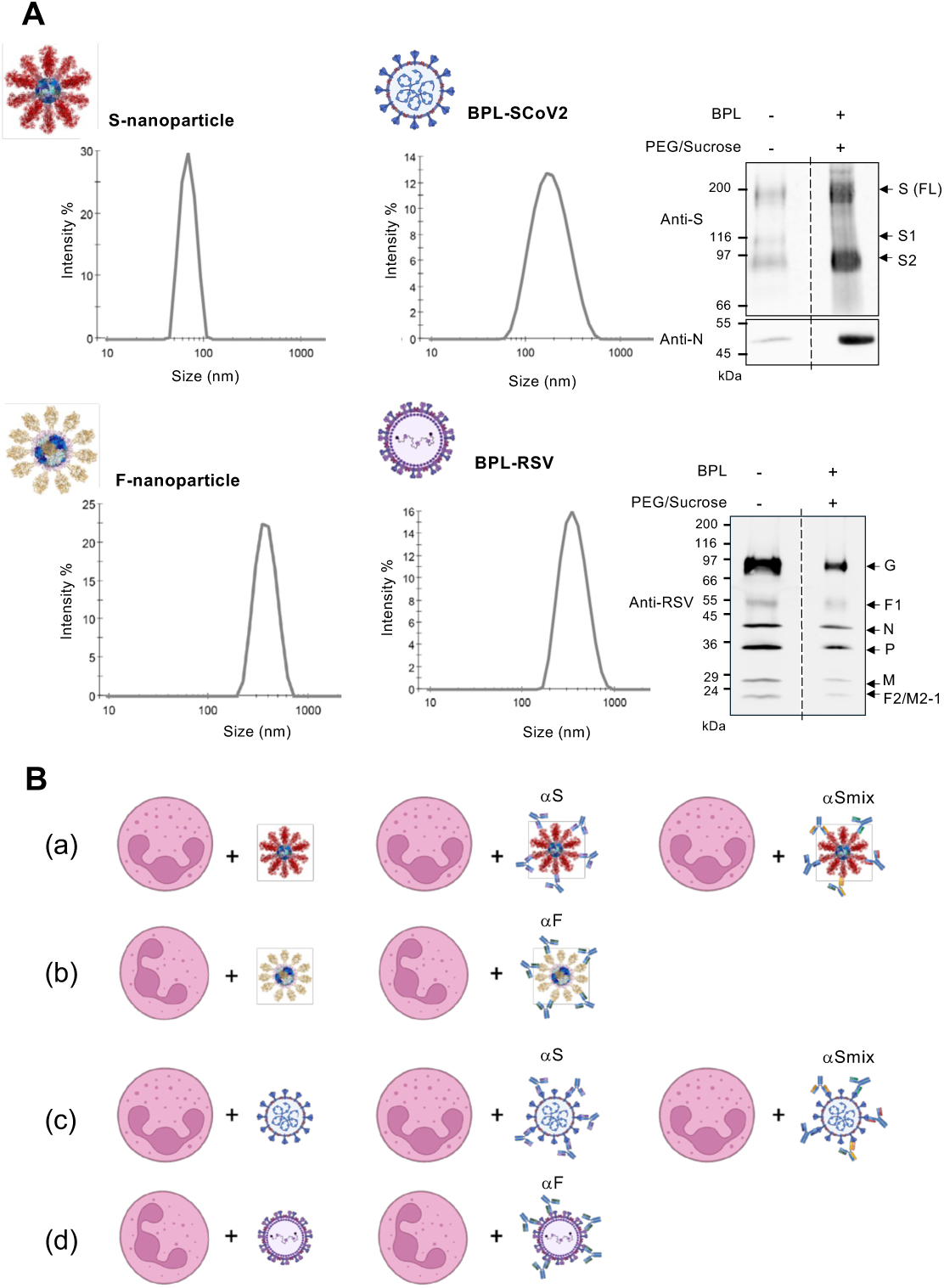
Experimental models. **(A)** Self-assembling nanoparticles displaying prefusion-stabilized trimeric SARS-CoV-2 Spike (S) or RSV Fusion (F) glycoproteins, as described in *Methods,* were analyzed by dynamic light scattering (DSL) for particle characterization. Purified β-propiolactone (BPL)-inactivated SARS-CoV-2 (SCoV2, Wuhan strain) and RSV A2 were assessed by immunoblot for viral proteins (S, N, or RSV antigens) and by dynamic light scattering (DSL). **(B)** *Ex vivo* models used in this study consisted of incubating human neutrophils with S-nanoparticles or BPL-SCoV2, either alone or pre-coated with anti-S antibodies—either a single monoclonal (αS) or a mix of five monoclonals (αSmix), as detailed in **Table 1**. Parallel experiments were conducted using F-nanoparticles or BPL-inactivated RSV A2.

Second, to assess the impact of the SARS-CoV-2 S protein within the structural context of a virus - and in a format relevant to inactivated virus vaccines - we used β-propiolactone (BPL)-inactivated, purified virus preparations. BPL-mediated inactivation is a widely used approach, notably employed in the CoronaVac (Sinovac) vaccine against SARS-CoV-2 [51]. Immunoblot analysis of purified, inactivated SARS-CoV-2 and RSV confirmed the presence of viral proteins (**Figure 1A**). Consistent with previous reports, BPL-inactivated SARS-CoV-2 virions exhibited a mix of prefusion and postfusion (mostly S2) peptides [52]. DSL analysis confirmed the uniformity and structural integrity of the virions, revealing single peaks at approximately 200 nm for SARS-CoV-2 and 300–400 nm for RSV (**Figure 1A)**. To assess the stability of our virus preparations, we analyzed freshly thawed virions as well as those stored at 4°C for various durations post-thaw using DSL measurements. In both cases, the preparations exhibited excellent stability for at least 48 hours (data not shown). Based on these findings, all subsequent experiments were conducted using thawed virion preparations stored at 4°C within this timeframe.

#### Impact of trimeric prefusion SARS-CoV-2 S on human neutrophils ex vivo

To assess the impact of the prefusion trimeric SARS-CoV-2 S glycoprotein on neutrophils, we utilized an *ex vivo* system in which primary human neutrophils were exposed to S-nanoparticles and for comparison to F-nanoparticles (**Figure 1B**). Neutrophil viability, surface expression of adhesion, degranulation, IgG interactions and complement activation and regulation markers, ROS production, NET formation, and cytokine levels were systematically measured.

Neutrophil viability was assessed after 24 h of exposure to increasing nanoparticle-to-neutrophil ratios, as described in *Methods*. Regardless of the S-nanoparticle concentration, approximately 55% of neutrophils remained viable, comparable to control conditions (**Figure 2A, B**). Similarly, F-nanoparticles protein had no discernible effect on neutrophil viability, further supporting the lack of cytotoxic impact (**Figure 2C**).

**Figure 2.**
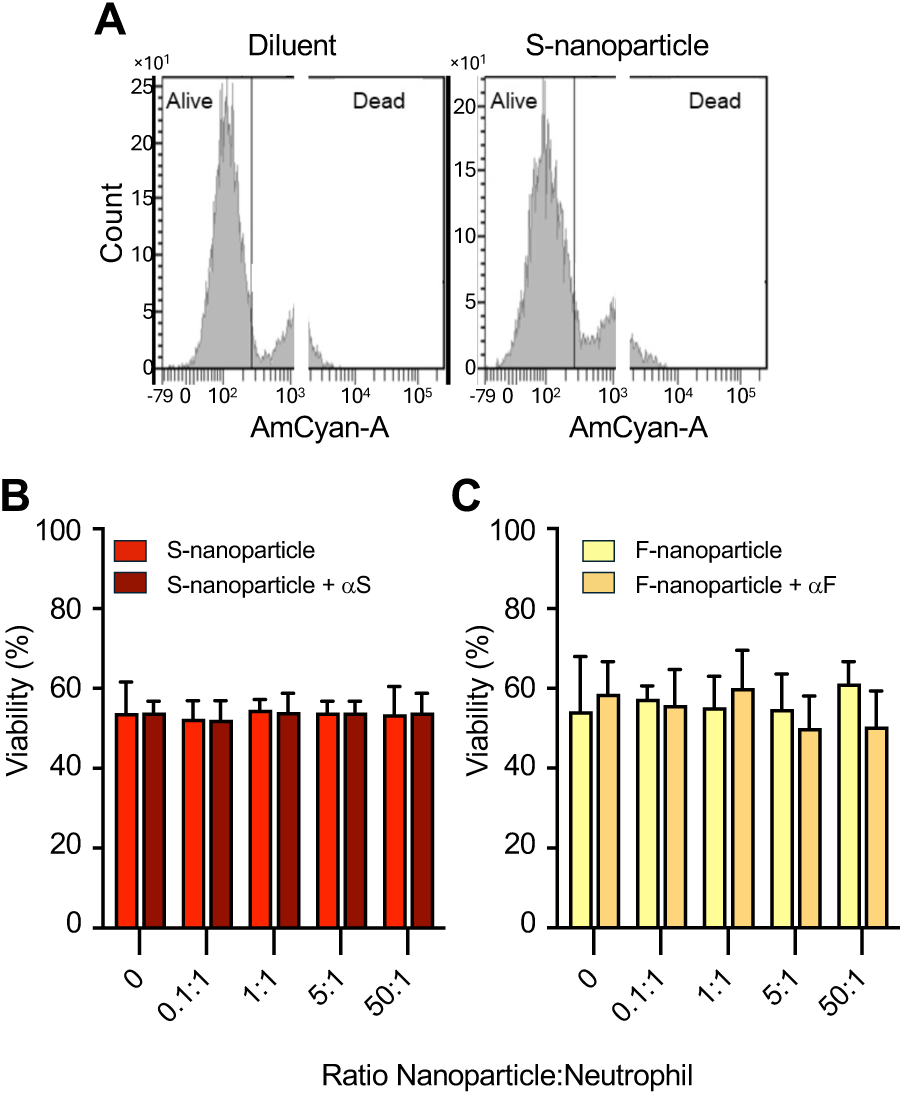
Effect of S-nanoparticles on neutrophil viability. **(A)** Human neutrophils were incubated for 24 h with S-nanoparticles or diluent control. Viability was assessed based on membrane integrity, as described in *Methods*. A representative experiment is shown. **(B-C)** Neutrophils were incubated for 24 h with the indicated S-**(B)** or F-nanoparticle **(C)**-to-neutrophil ratios, either alone or pre-coated with their respective antibodies (αS or αF, as in Figure 1). Data show mean ± SEM from three independent experiments, each using neutrophils from a different donor. Statistical analyses were performed using a two-way RM ANOVA.

Next, we examined a panel of neutrophil surface markers associated with adhesion (CD11b, CD15, CD62L), degranulation (CD63, CD66b), IgG interactions (CD16, CD32, CD64), and complement activation and regulation (CD46, CD55, CD59, CD93). Neutrophils were incubated with increasing nanoparticle-to-neutrophil ratios of S-nanoparticles for 30 min (**Figure 3**) or 3 h (**Supplementary Table 1**). Flow cytometry analysis, which we previously reported allows to detect variation in surface markers expression [43], revealed that S-nanoparticles had no significant impact on the expression of any of the quantified markers at any of the ratio tested (**Figure 3A-B**). These results are similar to surface markers expression observed when neutrophils were incubated with F-nanoparticles (**Figure 3B and Supplementary Table 1**).

**Figure 3.**
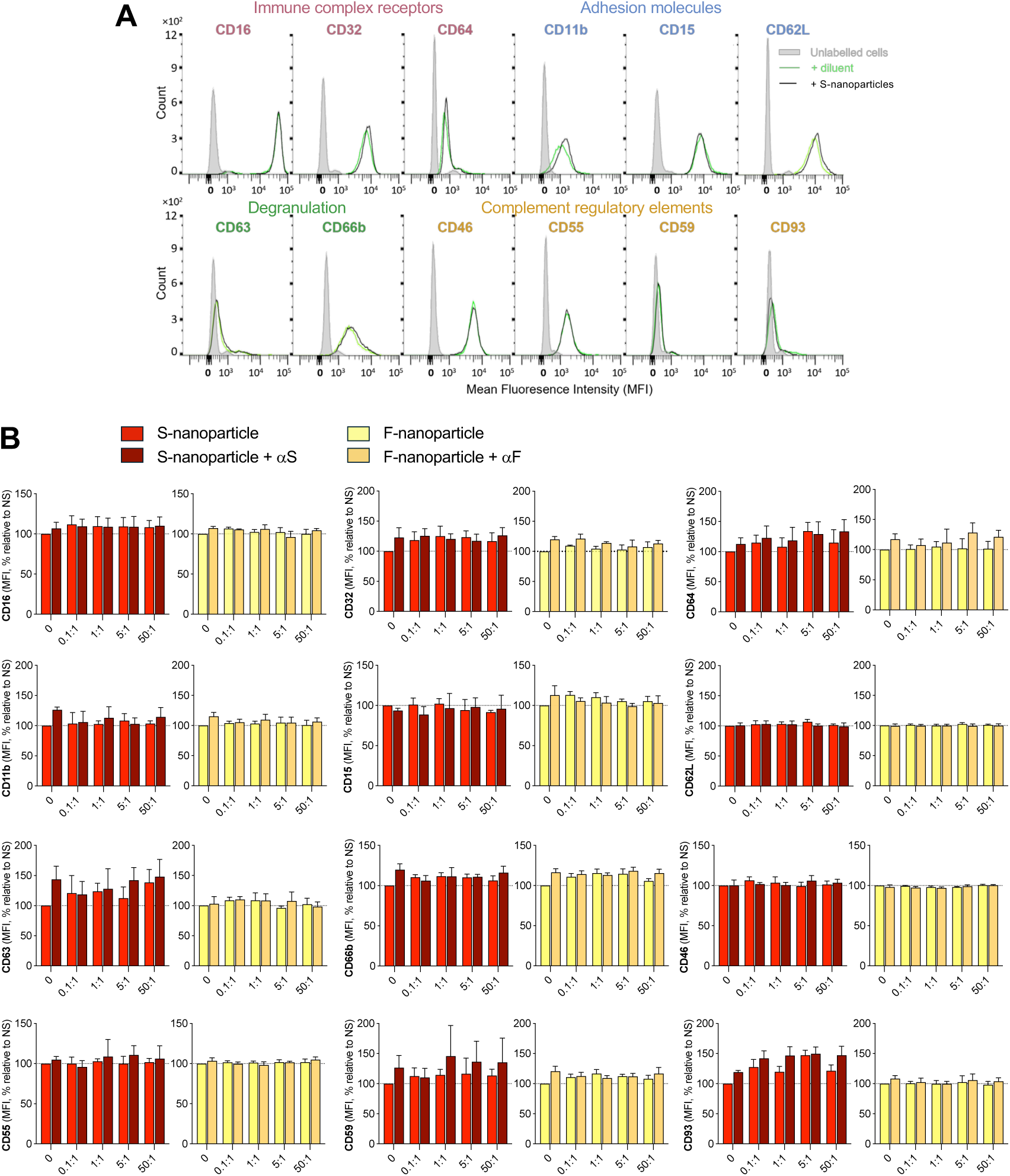
Impact of S-nanoparticles alone or pre-coated with anti-S antibodies on neutrophil surface marker expression. Neutrophils were incubated for 30 min or 3 h with the indicated S-nanoparticle-to-cell ratios, either alone or pre-coated with a monoclonal anti-S antibody (αS; see Figure 1). F-nanoparticles, with or without pre-coating with anti-F antibody (αF; see Figure 1), were used as comparators. Surface markers were stained with fluorophore-conjugated antibodies and analyzed by flow cytometry (**Table 2**). Markers were grouped by function: immune complex receptors (CD16, CD32, CD64), adhesion (CD11b, CD15, CD62L), degranulation (CD63, CD66b), and complement regulation (CD46, CD55, CD59, CD93). (A) A representative experiment using a S-nanoparticle-to-cell ratio of 50:1 incubated for 30 min is shown. (B) Quantified data from 30 min incubations are shown as mean ± SEM (n = 3 donors). Surface expression is reported as percent change in mean fluorescence intensity (MFI) relative to non-stimulated (NS) cells. Full results, including 3 h data, are presented in **Supplementary Table 1**.

As key players in the innate immune response, neutrophils generate ROS and NETs in response to various stimuli. Elevated ROS levels correlate with neutrophil counts in patients with severe COVID-19 [53], while excessive NETosis has been associated with disease severity [14–16]. However, the role of the prefusion trimeric S glycoprotein in these processes remains unknown. To assess whether S-nanoparticles could induce ROS production or modulate neutrophil ROS responses to established stimuli, we measured ROS levels following S-nanoparticle exposure in the presence or absence of heat-aggregated (HA)-IgGs, a model for immune complexes [54]. S-nanoparticles had no discernible impact on ROS production, either alone or in combination with HA-IgGs, a finding that mirrored results observed with F-nanoparticles (**Figure 4A-B**). We next investigated whether S-nanoparticles could directly induce NET formation. Unlike PMA, a well-established NETosis inducer, S-nanoparticles had no significant effect on NET production. F-nanoparticles similarly failed to induce NETosis (**Figure 4C**).

**Figure 4.**
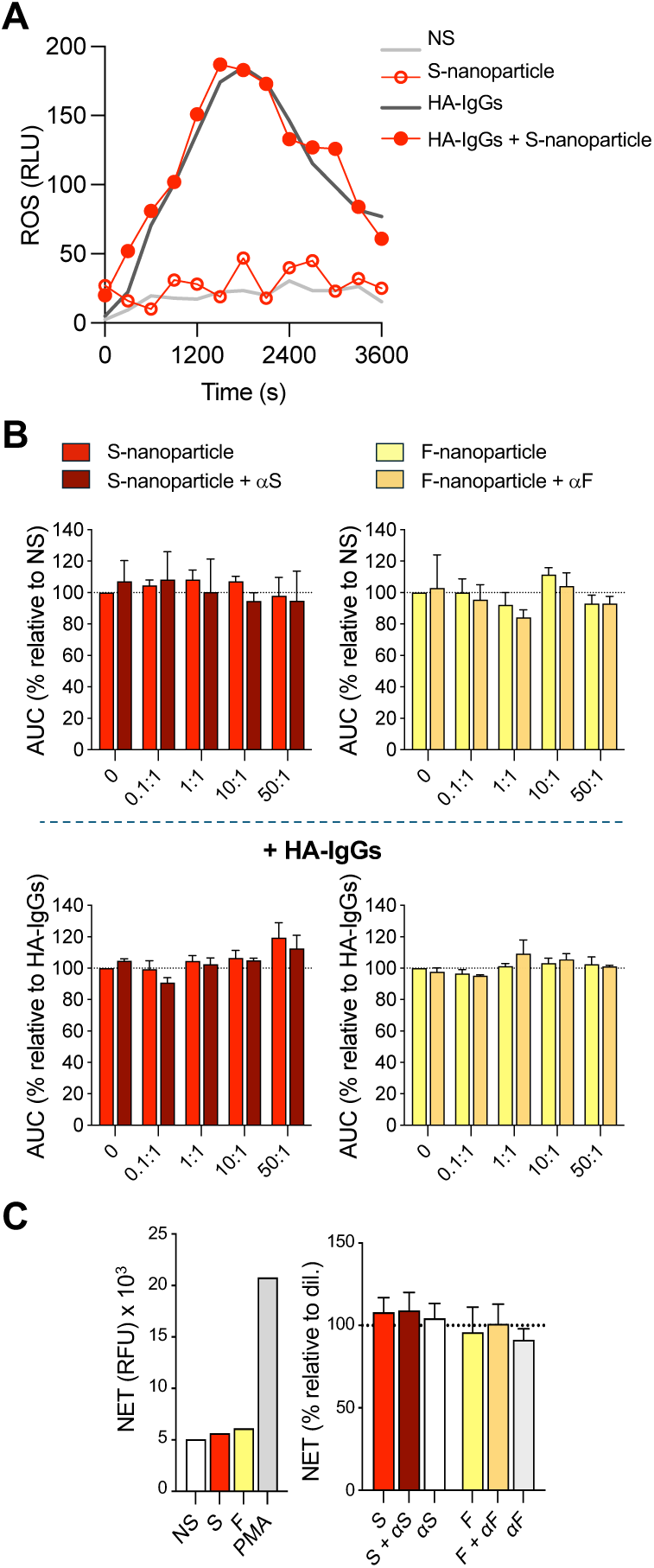
Effect of S-nanoparticles on neutrophil ROS production and NET formation. Human neutrophils were left unstimulated (NS) or incubated with the indicated S-nanoparticle-to-cell ratios, either alone or in the presence of 1 mg/mL heat-aggregated immunoglobulins (HA-IgGs). S-nanoparticles were used either uncoated or pre-coated with anti-S antibody (αS; Figure 1). F-nanoparticles, with or without anti-F antibody (αF, Figure 1), were included as comparators. (**A–B**) ROS production was monitored over 60 min by luminol-based chemiluminescence. (**A**) A representative experiment using a nanoparticle-to-neutrophil ratio of 50:1 is shown; results are expressed as relative luminescence units (RLU). (**B**) ROS levels are quantified as area under the curve (AUC) and expressed as percentages relative to NS or HA-IgG-stimulated cells, as indicated. (C) NET formation was measured after 4 h of incubation at a nanoparticle-to-neutrophil ratio of 50:1 using SYTOX green staining, as described in *Methods*. PMA (10 nM) was used as a positive control. Left: representative experiment, expressed as relative fluorescence units (RFU). Right: NET production expressed as percentage relative to NS. Data in (**B**) and (**C**) are shown as mean ± SEM from n = 3 independent experiments with neutrophils from different donors.

Cytokine storm is a hallmark of COVID-19 complications [3], and neutrophils are thought to contribute to the excessive inflammation. To investigate the inflammatory gene response, we exposed neutrophils to S-nanoparticles for 60 min and measured the expression of genes encoding *COX-2*, *CCL3* (MIP-1α), *CXCL8* (IL-8), and *IFNB* (IFNβ). An increasing trend, although not reaching statistical significance, was observed for *COX-2*, *CCL3*, and *IFNB* mRNA (**Figure 5A**). We also examined cytokine secretion by measuring IL-1α, IL-1β, IL-6, CXCL8 (IL-8), TNF, IFNα, CCL3 (MIP-1α), and CCL2 (MCP-1) in the supernatants from neutrophils exposed to nanoparticles for 4 h. Of these, only IL-8 was detected. Neither S- nor F-nanoparticles affected IL-8 levels of cytokines (**Figure 5B**).

**Figure 5.**
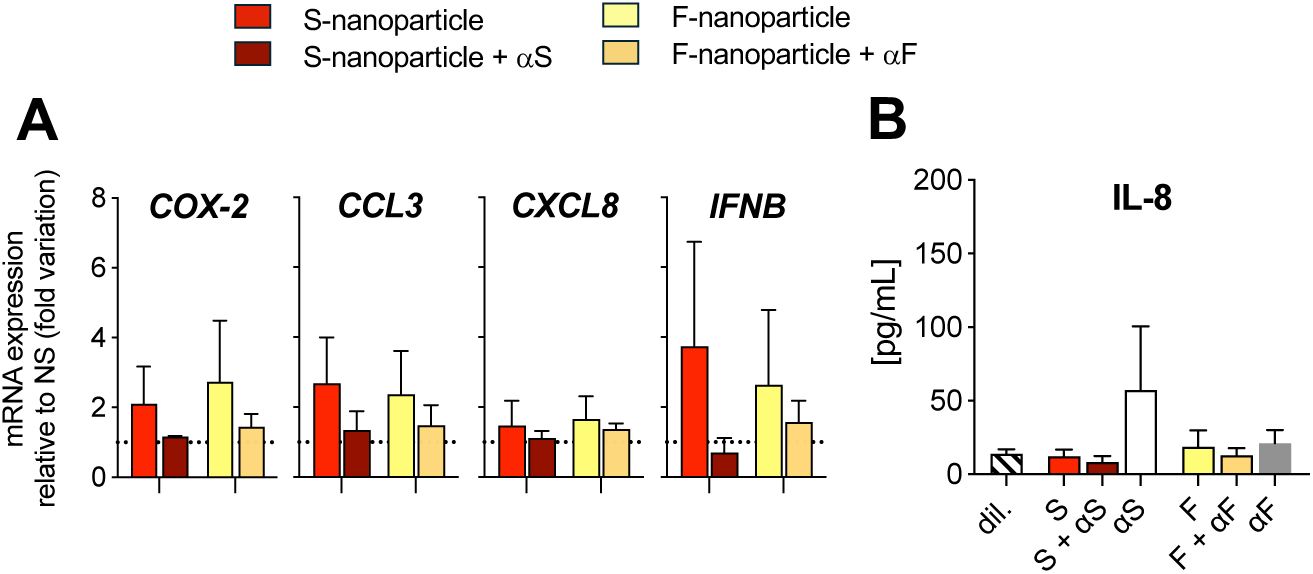
Effect of S-nanoparticles on inflammatory gene expression and cytokine release. Human neutrophils were incubated for 60 min with S- or F-nanoparticles at a nanoparticle-to-cell ratio of 50:1, either uncoated or pre-coated with their respective antibodies (Figure 1). (**A**) mRNA levels of the indicated genes were quantified by RT-qPCR, normalized to GAPDH, and expressed as fold change relative to non-stimulated cells (NS). (**B**) IL-8 levels in the supernatant were quantified using multiplexed bead-based immunoassay. All data represent mean ± SEM from n = 3 independent experiments using neutrophils from different donors.

#### Inactivated SARS-CoV-2 has minimal impact on neutrophil responses

Results obtained with S-nanoparticles provided no evidence that the trimeric prefusion-stabilized S protein significantly influences neutrophil responses. To investigate whether the S protein in the context of a viral structure - relevant to inactivated virus vaccines - elicits different effects, we examined the impact of the purified BPL-inactivated SARS-CoV-2 in the *ex vivo* system with primary human neutrophils (**Figure 1B**). For comparison, we also used BPL-inactivated RSV. Neutrophil viability remained unaffected following 24 h exposure to increasing ratios of inactivated SARS-CoV-2 or RSV to neutrophils (**Figure 6A**). Expression of most surface markers associated with degranulation, IgG interactions, and complement activation and regulation was largely unchanged after 30 min or 3 h exposure to inactivated SARS-CoV-2. (Full results are presented in **Supplementary Table 2**). However, exposure to inactivated SARS-CoV-2 led to an upregulation of CD11b—and to a lesser extent, CD62L—both of which are associated with cell adhesion. This effect was observed after 3 h and 30 min of exposure, respectively. Induction of CD62L was also observed after exposure to inactivated RSV for 3 h. An increasing, but modest, trend of CD32 expression, associated with IgG interactions, was also observed after 3 h of exposure to inactivated SARS-CoV-2 (**Figure 6B**). Inactivated SARS-CoV-2 had no influence on ROS production **(Figure 6C**) in the absence or presence of HA-IgGs, NET formation (**Figure 6D**), inflammatory gene expression (**Figure 6E**), or IL-8 release (**Figure 6F**) at any of the virus particle-to-neutrophil ratio tested. These results are similar to neutrophil treatment with inactivated RSV. Collectively, these findings reinforce the conclusion that the SARS-CoV-2 S protein, even in the context of an inactivated virus, has little to no effect on the tested neutrophil responses.

**Figure 6.**
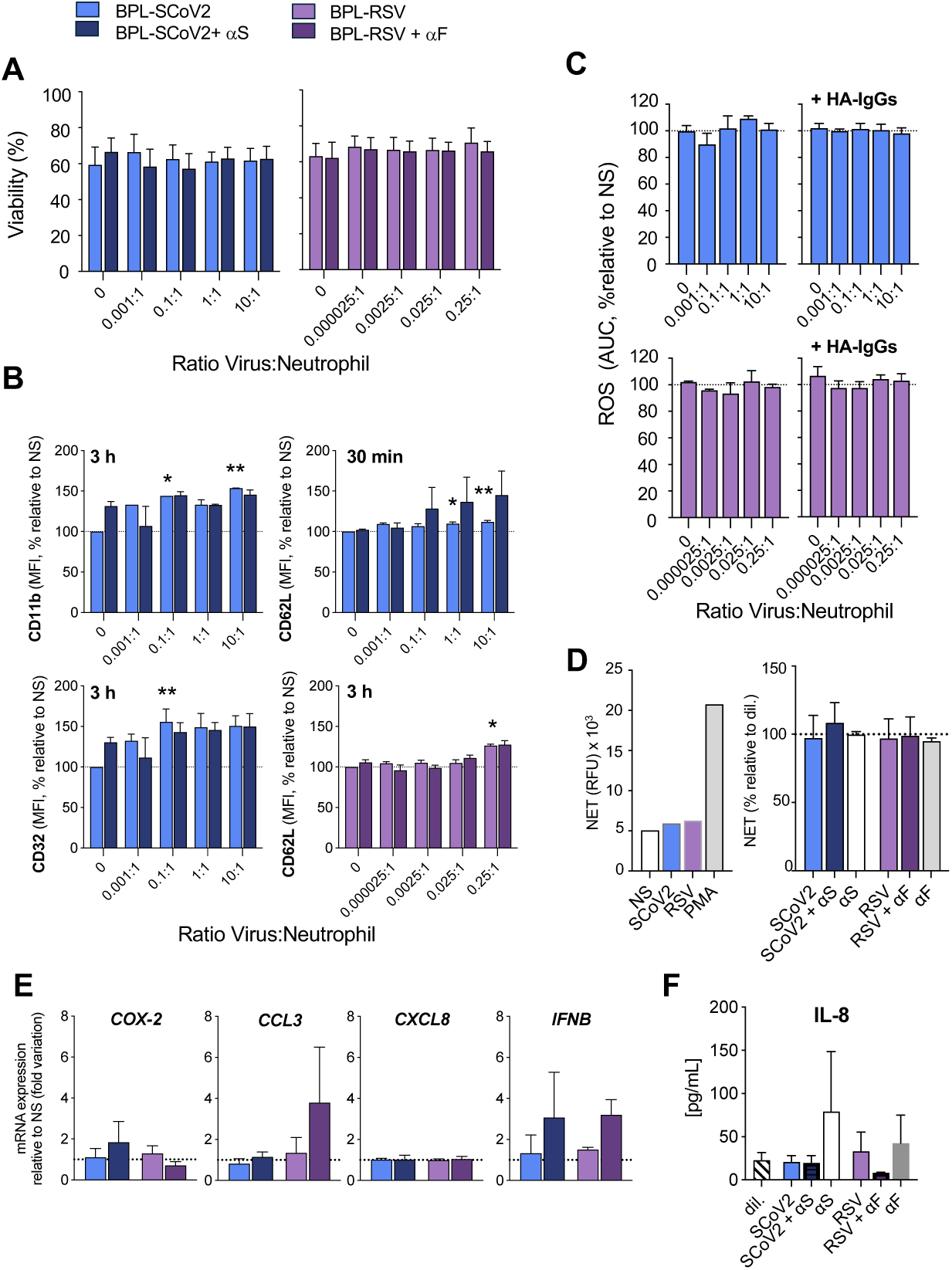
Effect of inactivated SARS-CoV-2 on neutrophil responses. Human neutrophils were incubated with the indicated ratios of BPL-inactivated SARS-CoV-2 (SCoV2), either uncoated or pre-coated with anti-S antibody (αS). BPL-inactivated RSV, with or without anti-F antibody (αF), was used as a comparator (Figure 1). (**A**) Neutrophil viability was assessed after 24 h, as described in Figure 2. (**B**) Surface expression of markers related to immune complex receptors (CD16, CD32, CD64), adhesion (CD11b, CD15, CD62L), degranulation (CD63, CD66b), and complement regulation (CD46, CD55, CD59, CD93) was measured by flow cytometry after 30 min and 3 h of incubation, as in Figure 3. Shown are markers with observed changes; full results are available in **Supplementary Table 2**. Data are expressed as percent change in mean fluorescence intensity (MFI) relative to non-stimulated (NS) cells. (**C**) ROS production was measured in the absence or presence of HA-IgGs, as in Figure 4. Results are expressed as area under the curve (AUC), relative to NS or HA-IgG-stimulated cells. (**D**) NET formation was quantified after 4 h, as in Figure 4. Left: A representative experiment using a virus-to-neutrophil ratio of 10:1 is shown. PMA (10 nM) was used as a positive control; results are expressed as relative luminescence units (RLU). Right: NET production expressed as percentage relative to NS. (**E**) mRNA levels of selected inflammatory genes were measured by RT-qPCR, as described in Figure 5. IL-8 levels in the supernatant were quantified using multiplexed bead-based immunoassay. All data represent mean ± SEM from n = 3 independent experiments using neutrophils from different donors. * *P*<0.05; ** *P*<0.01.

#### Antibody binding to Spike does not alter its effect on neutrophil responses

A diverse array of anti-S antibodies is generated following SARS-CoV-2 infection or vaccination [55]. Consequently, upon subsequent encounters, the S glycoprotein is likely to be antibody-bound, facilitating recognition by phagocytes, including neutrophils. To assess the effect of antibody-bound S glycoprotein on neutrophils in the *ex vivo* system, we first pre-coated S-nanoparticles with a single neutralizing monoclonal anti-S antibody (αS, **Table 1 and Figure 1B**). This antibody, derived from B cells specific for the SARS-CoV-2 S glycoprotein, was isolated from a COVID-19-infected individual and targets the receptor-binding domain (RBD; residues 319–591), effectively blocking the RBD-ACE2 interaction [56]. For comparison, we pre-coated F-nanoparticles with Palivizumab, a clinically approved humanized monoclonal antibody (αF; **Table 1 and Figure 1B**) used to prevent severe lower respiratory tract infections (LRTI) caused by RSV. Palivizumab prevents virus entry by inhibiting viral-host membrane fusion and may also suppress cell-to-cell transmission by blocking syncytia formation in respiratory epithelial cells. As a result, it reduces RSV virulence and the risk of RSV-related LRTI [57]. In this experimental setting, antibody-bound S- and F-nanoparticles had no significant effect on neutrophil viability (**Figure 2B**), surface marker expression (**Figure 3B and Supplementary Table 1**), ROS production (**Figure 4B**), NET formation (**Figure 4C**), inflammatory gene expression (**Figure 5A**), or cytokine secretion (**Figure 5B**). Similarly, neutrophil viability (**Figure 6A**), ROS production (**Figure 6C)**, NETosis (**Figure 6D**), and IL-8 release were unaffected by inactivated SARS-CoV-2 pre-coated with αS. Amongst surface marker expression, a non-significant trend toward increased CD62L expression was observed in response to antibody-coated SARS-CoV-2 (**Figure 6B**). This trend was not observed with αF-coated inactivated RSV (**Figure 6B**). Similarly, antibody-coated inactivated SARS-CoV-2 and RSV were associated with a non-significant trend toward increased *IFNB* gene expression (**Figure 6E**).

Since natural and vaccine-induced immune responses generate polyclonal antibodies against multiple S epitopes, we next examined whether a diverse antibody mix altered neutrophil responses differently than a single neutralizing antibody. This approach provides a more physiologically relevant model of immune complex formation and FcγR-mediated neutrophil activation. S-nanoparticles or BPL-inactivated SARS-CoV-2 were pre-coated with a mix of five anti-S antibodies (αSmix; **Table 1 and Figure 1B**) before exposure to neutrophils. Coating with the αSmix did not impact neutrophil viability, surface marker expression, ROS and NET production compared to coating with αS (**Figure 7 and Supplementary Table 3**). However, a non-significant trend toward increased *COX2* and *CXCL8* gene expression (**Figure 7D**) and IL-8 release (**Figure 7E**) was noticed.

**Figure 7.**
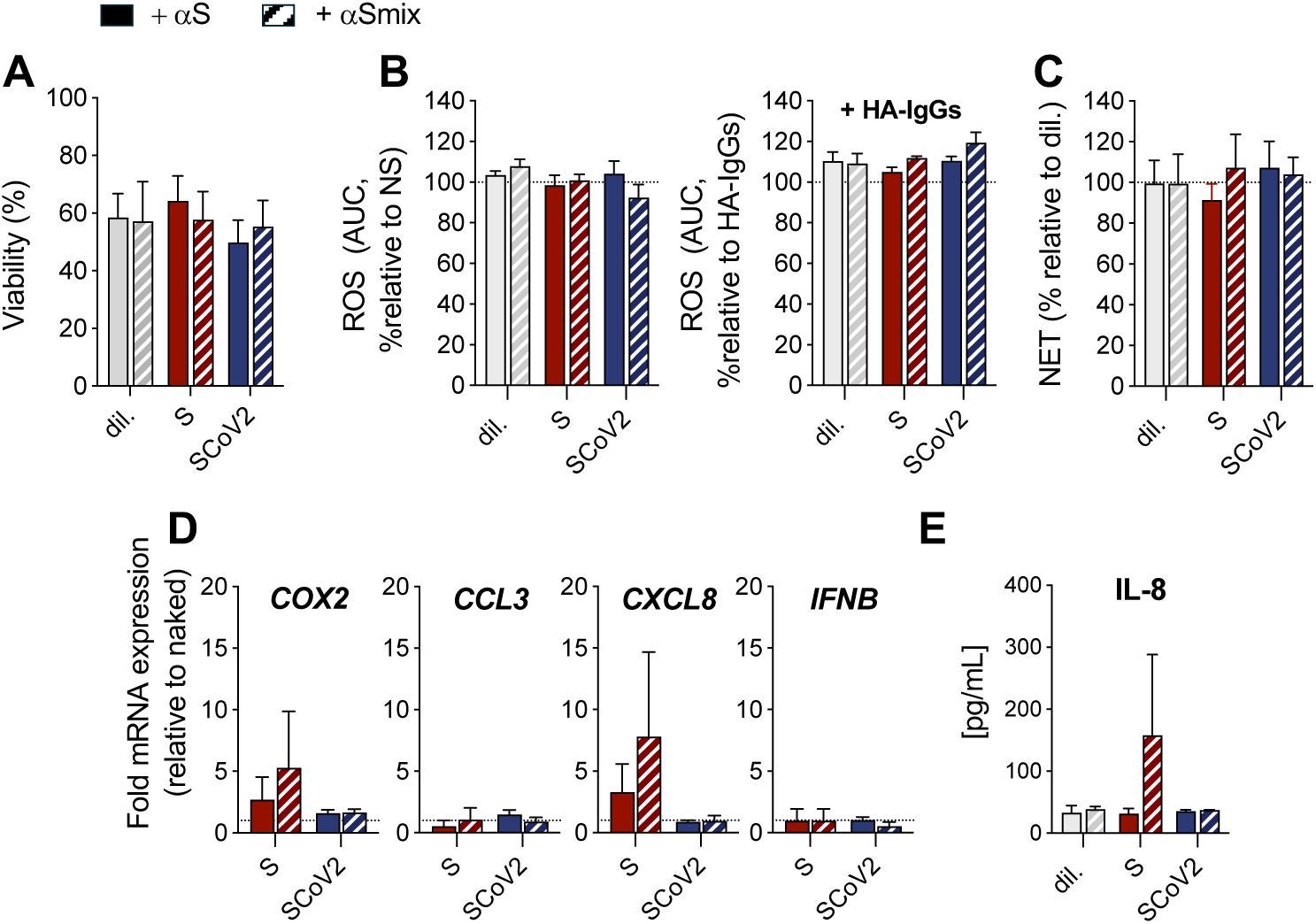
Comparison of single vs multiple anti-Spike antibodies for nanoparticle and virus pre-coating on neutrophil responses. Neutrophils were incubated with diluent (dil), S-nanoparticles, or BPL-inactivated SARS-CoV-2 (SCoV2), either pre-coated with a single monoclonal anti-S antibody (αS) or with a mixture of five monoclonal anti-S antibodies (αSmix), as described in Figure 1. Neutrophil viability **(A)**, ROS production **(B)**, NET formation **(C)**, inflammatory gene expression **(D)**, and IL-8 release **(E)** were assessed and analyzed as described in Figures 2–5. Surface marker expression data are presented in **Supplementary Table 3**. All results are shown as mean ± SEM from n = 3 independent experiments, each performed with neutrophils from a different donor.

Collectively, these findings indicate that antibody binding does not significantly alter the neutrophil response to the S protein, whether using a single neutralizing antibody or a diverse antibody mix. These results suggest that FcγR engagement by Spike-containing immune complexes is insufficient to drive substantial neutrophil activation under these conditions.

## Discussion

The SARS-CoV-2 S protein is essential for viral entry and is the principal target of neutralizing antibodies, making it a key component in COVID-19 vaccines. Understanding how the S protein interacts with innate immune cells, especially neutrophils, is critical given their dual role in antiviral defense and inflammatory pathology. In this study, we evaluated whether the prefusion-stabilized trimeric S protein directly induces neutrophil responses *ex vivo*. Using two biologically relevant models—nanoparticle-displayed S and BPL-inactivated SARS-CoV-2—we found no evidence that S alone, in these contexts, elicits robust neutrophil responses.

Our models were selected to reflect relevant vaccine contexts. The S-nanoparticles mimic the multivalent display of prefusion S found on intact virions, recombinant protein nanoparticle vaccines (e.g., Novavax), and membrane-anchored S expressed on host cells following mRNA vaccination. The BPL-inactivated virus, similar to the CoronaVac vaccine, presents a mixture of prefusion and postfusion Spike conformations within the native virion structure, as previously reported [52]. These models therefore encompass different physical presentations and contexts of the S protein relevant to both infection and vaccination.

Most previous studies reporting S-mediated neutrophil activation employed soluble recombinant S protein, which differs markedly from the multivalent, trimeric, and membrane-bound or particle-anchored forms used here. This difference in presentation is significant, as the conformation, oligomerization state, and valency of the S protein are likely to influence receptor engagement and downstream immune activation. Supporting the relevance of our model, the same S-nanoparticles used in this study have previously elicited potent neutralizing responses—25-fold higher than trimeric soluble S protein—in a mouse immunization model at low antigen doses [35]. Despite this immunogenicity, we found that S-nanoparticles induced little to no neutrophil responses under the tested conditions. Across a range of nanoparticle-to-neutrophil ratios, we observed no significant modulation of surface markers reflecting adhesion, degranulation, IgG interactions and complement activation and regulation, ROS production, NETosis, inflammatory gene expression or cytokine release. In line with our findings, prior reports have produced conflicting evidence regarding neutrophil activation by recombinant S. One study found that full-length recombinant S, but not S1 or S2 subunits, could induce NETs in a ROS-independent manner at high S:neutrophil ratios [58, 59], while another observed increased ROS and IL-8/IL1-Ra in response to recombinant S1 protein [60]. In contrast, Ait-Belkacem et al. reported that trimeric prefusion S induced only minimal granulocyte responses in whole blood, with no significant changes in any neutrophil activation markers and only a trend toward higher CD64 expression [61]. Additionally, Veras et al. demonstrated that live, replicating SARS-CoV-2 is required for NETosis induction, suggesting that S alone may be insufficient [14]. This aligns with our observation that BPL-inactivated SARS-CoV-2 failed to induce detectable neutrophil activation.

Glycosylation is another critical determinant of S immunogenicity and receptor interactions. The glycan composition of S protein can vary significantly between expression systems, affecting how immune cells recognize and respond to it. The SARS-CoV-2 S protein contains over 20 N-linked glycosylation sites, which influence epitope masking and structural stability [62]. Studies show that removing the glycan shield enhances immunogenicity and protection [63]. The impact of BPL inactivation on S glycosylation is not fully characterized, though the efficacy of CoronaVac suggests preserved recognition by immune cells. Similarly, S-nanoparticles were produced in a mammalian expression system designed to generate glycosylated proteins with appropriate immunogenic features. While subtle glycosylation differences may exist, the potent immune responses observed in prior mouse studies argue against a major impairment in antigenicity or neutrophil recognition [35].

In this study, we evaluated S from the Wuhan strain in the BPL-inactivated virus and the D614G variant in the S-nanoparticles model. Since S mutations can modulate ACE2 binding, immune escape, and antigenicity, it remains possible that variant-specific differences influence neutrophil responses. For instance, the Omicron variant carries mutations that alter both antibody binding and innate immune engagement. Therefore, we cannot exclude that different results would be obtained with S protein from different variants.

Importantly, in real-world settings, neutrophils encounter the S protein in the context of inflammation, cytokine gradients, and immune complexes—none of which are fully captured in our *ex vivo* system. We did model one such context by pre-coating S-nanoparticles with anti-S antibodies, mimicking the formation of immune complexes following infection or vaccination. Even under these conditions, we found no evidence of enhanced neutrophil responses via Fcγ receptor engagement, either with a neutralizing monoclonal antibody or a mixture of monoclonals targeting diverse S epitopes. These data suggest that immune complex formation alone may be insufficient to trigger neutrophil responses in the absence of additional inflammatory or danger signals.

## Acknowledgments

This work was funded by the Support Program for Research and Innovation Organizations, from the Ministre de l’économie et de l’innovation du Québec (MEI, Grant no 2020-2021-COVID-19-PSOv2a) to MP and NG, funds from the Fondation du CHUM to NG, and a grant from the Fondation du CHU de Québec to MP (grant no. 4004). The authors thank Dr. Simon Grandjean Lapierre and Floriane Point (CRCHUM, Montreal, Canada) for SARS-CoV-2 sequencing, Dr. Samira Mubareka (Sunnybrook Research Institute, Toronto, Canada) for the SARS-CoV-2/SB2 isolate and Dr. A. Banerjee (VIDO, Saskatoon University) for advice with SARS-CoV-2 culture. The authors also thank Drs. A. Jureka (Center for Microbial Pathogenesis, Atlanta, USA) and C. Basler (Icahn School of Medicine at Mount Sinaï, New-York, USA) for advice with SARS-CoV-2 inactivation. Figure 1 was

## Authors’ Contributions

MP and NG contributed to the design of the study. AF, SH, EC, CL, AP and CG contributed to data collection. BZ and PDK produced and validated nanoparticles. MP and NG conducted data analysis and interpretation. MP and NG drafted the first version of the article. AF, SH, EC, CL, AP, NC, CG, BZ, PDK, MP and NG critically revised the article and approved the final version for submission for publication. Figure 1 created in BioRender. Grandvaux, N. (2025) https://BioRender.com/3mmudj8

