## Supplemental Table 1 for "No Evidence of Direct Activation of Human Neutrophil Responses by Multivalent Prefusion Trimeric SARS-CoV-2 Spike Protein *ex vivo*"

**A)**

| | | | S | | | S + $\alpha$S | | | F | | | F + $\alpha$F | | |
|---|---|---|---|---|---|---|---|---|---|---|---|---|---|---|
| | | Ratio | % | SEM | *P* | % | SEM | *P* | % | SEM | *P* | % | SEM | *P* |
| **CD16** | **30 min** | 0 | 100.00 | 0.00 | n.s. | 106.88 | 7.57 | n.s. | 100.00 | 0.00 | n.s. | 107.12 | 2.20 | n.s. |
| | | 0.1:1 | 111.93 | 10.79 | n.s. | 109.50 | 9.03 | n.s. | 106.66 | 1.73 | n.s. | 105.55 | 0.73 | n.s. |
| | | 1:1 | 109.69 | 11.77 | n.s. | 108.96 | 10.74 | n.s. | 102.49 | 2.99 | n.s. | 106.20 | 5.03 | n.s. |
| | | 5:1 | 109.26 | 11.18 | n.s. | 108.87 | 12.87 | n.s. | 102.31 | 5.43 | n.s. | 96.26 | 6.87 | n.s. |
| | | 50:1 | 108.42 | 8.25 | n.s. | 110.19 | 11.02 | n.s. | 99.98 | 5.71 | n.s. | 104.44 | 2.34 | n.s. |
| | **3 hrs** | 0 | 100.00 | 0.00 | n.s. | 108.04 | 3.83 | n.s. | 100.00 | 0.00 | n.s. | 99.45 | 6.63 | n.s. |
| | | 0.1:1 | 106.13 | 2.70 | n.s. | 99.00 | 12.89 | n.s. | 101.80 | 1.09 | n.s. | 108.13 | 5.14 | n.s. |
| | | 1:1 | 96.12 | 10.31 | n.s. | 113.84 | 4.61 | n.s. | 105.43 | 2.03 | n.s. | 107.18 | 7.08 | n.s. |
| | | 5:1 | 99.97 | 14.58 | n.s. | 118.27 | 2.23 | n.s. | 105.56 | 5.48 | n.s. | 104.76 | 7.09 | n.s. |
| | | 50:1 | 94.24 | 15.80 | n.s. | 108.70 | 2.02 | n.s. | 108.22 | 8.22 | n.s. | 108.33 | 8.20 | n.s. |
| **CD32** | **30 min** | 0 | 100.00 | 0.00 | n.s. | 122.91 | 15.99 | n.s. | 100.00 | 0.00 | n.s. | 119.94 | 4.72 | n.s. |
| | | 0.1:1 | 118.44 | 13.77 | n.s. | 125.56 | 12.13 | n.s. | 109.35 | 1.17 | n.s. | 121.02 | 7.59 | n.s. |
| | | 1:1 | 125.18 | 16.50 | n.s. | 120.55 | 8.39 | n.s. | 104.26 | 4.03 | n.s. | 114.17 | 2.15 | n.s. |
| | | 5:1 | 123.35 | 10.42 | n.s. | 117.21 | 10.98 | n.s. | 102.99 | 7.79 | n.s. | 108.08 | 10.68 | n.s. |
| | | 50:1 | 116.54 | 13.97 | n.s. | 126.36 | 12.99 | n.s. | 107.07 | 8.72 | n.s. | 113.08 | 5.32 | n.s. |
| | **3 hrs** | 0 | 100.00 | 0.00 | n.s. | 103.95 | 6.88 | n.s. | 100.00 | 0.00 | n.s. | 101.834 | 6.0888 | n.s. |
| | | 0.1:1 | 104.80 | 9.14 | n.s. | 89.81 | 19.07 | n.s. | 108.81 | 6.98 | n.s. | 100.928 | 2.7957 | n.s. |
| | | 1:1 | 88.07 | 14.14 | n.s. | 104.11 | 8.49 | n.s. | 98.56 | 5.96 | n.s. | 100.654 | 5.4654 | n.s. |
| | | 5:1 | 93.09 | 20.96 | n.s. | 102.54 | 10.38 | n.s. | 105.90 | 5.55 | n.s. | 97.749 | 1.8982 | n.s. |
| | | 50:1 | 91.98 | 22.36 | n.s. | 103.34 | 8.85 | n.s. | 107.42 | 4.68 | n.s. | 113.074 | 13.578 | n.s. |
| **CD64** | **30 min** | 0 | 100.00 | 0.00 | n.s. | 112.51 | 10.51 | n.s. | 100.00 | 0.00 | n.s. | 117.33 | 8.76 | n.s. |
| | | 0.1:1 | 114.83 | 12.41 | n.s. | 122.71 | 19.85 | n.s. | 101.08 | 6.94 | n.s. | 107.52 | 10.22 | n.s. |
| | | 1:1 | 108.05 | 14.96 | n.s. | 118.57 | 21.74 | n.s. | 105.02 | 8.37 | n.s. | 111.53 | 22.60 | n.s. |
| | | 5:1 | 134.05 | 14.40 | n.s. | 129.10 | 20.41 | n.s. | 102.10 | 15.96 | n.s. | 128.49 | 15.92 | n.s. |
| | | 50:1 | 114.86 | 21.53 | n.s. | 133.61 | 19.51 | n.s. | 101.67 | 12.09 | n.s. | 121.21 | 10.85 | n.s. |
| | **3 hrs** | 0 | 100.00 | 0.00 | n.s. | 105.32 | 4.73 | n.s. | 100.00 | 0.00 | n.s. | 104.95 | 7.40 | n.s. |
| | | 0.1:1 | 103.17 | 4.15 | n.s. | 102.51 | 9.87 | n.s. | 100.16 | 5.91 | n.s. | 105.07 | 2.80 | n.s. |
| | | 1:1 | 106.05 | 9.94 | n.s. | 97.63 | 4.97 | n.s. | 100.51 | 5.57 | n.s. | 112.25 | 7.81 | n.s. |
| | | 5:1 | 106.18 | 13.93 | n.s. | 113.22 | 9.46 | n.s. | 108.37 | 4.44 | n.s. | 115.22 | 5.24 | n.s. |
| | | 50:1 | 101.89 | 7.59 | n.s. | 102.64 | 4.35 | n.s. | 100.03 | 5.55 | n.s. | 117.09 | 5.95 | n.s. |

**B)**

| | | | S | | | S + $\alpha$S | | | F | | | F + $\alpha$F | | |
|---|---|---|---|---|---|---|---|---|---|---|---|---|---|---|
| | | Ratio | % | SEM | *P* | % | SEM | *P* | % | SEM | *P* | % | SEM | *P* |
| **CD11b** | **30 min** | 0 | 100.00 | 0.00 | n.s. | 126.38 | 4.39 | n.s. | 100 | 0 | n.s. | 115.375 | 6.5644 | n.s. |
| | | 0.1:1 | 103.46 | 18.16 | n.s. | 106.05 | 17.69 | n.s. | 104.233 | 3.124 | n.s. | 105.656 | 4.8279 | n.s. |
| | | 1:1 | 102.58 | 5.37 | n.s. | 112.97 | 18.21 | n.s. | 103.299 | 4.0964 | n.s. | 109.635 | 9.2132 | n.s. |
| | | 5:1 | 108.46 | 11.65 | n.s. | 103.17 | 9.83 | n.s. | 104.854 | 9.0962 | n.s. | 104.808 | 9.1657 | n.s. |
| | | 50:1 | 103.55 | 4.37 | n.s. | 114.51 | 15.59 | n.s. | 100.693 | 8.0376 | n.s. | 106.482 | 6.4187 | n.s. |
| | **3 hrs** | 0 | 100.00 | 0.00 | n.s. | 116.22 | 3.82 | n.s. | 100 | 0 | n.s. | 98.6023 | 9.1201 | n.s. |
| | | 0.1:1 | 109.55 | 0.80 | n.s. | 117.81 | 2.53 | n.s. | 101.174 | 0.968 | n.s. | 102.737 | 4.0444 | n.s. |
| | | 1:1 | 108.70 | 3.16 | n.s. | 118.20 | 9.25 | n.s. | 96.2682 | 3.7248 | n.s. | 105.537 | 6.9425 | n.s. |
| | | 5:1 | 119.75 | 11.24 | n.s. | 135.73 | 0.54 | n.s. | 107.154 | 5.4448 | n.s. | 105.986 | 7.3046 | n.s. |
| | | 50:1 | 107.16 | 8.77 | n.s. | 106.27 | 6.60 | n.s. | 104.183 | 5.8682 | n.s. | 109.399 | 12.137 | n.s. |
| **CD15** | **30 min** | 0 | 100.00 | 0.00 | n.s. | 93.65 | 2.84 | n.s. | 100.00 | 0.00 | n.s. | 112.89 | 11.77 | n.s. |
| | | 0.1:1 | 101.09 | 8.08 | n.s. | 88.68 | 9.93 | n.s. | 112.98 | 4.48 | n.s. | 105.64 | 3.80 | n.s. |
| | | 1:1 | 102.22 | 6.41 | n.s. | 96.63 | 18.41 | n.s. | 110.29 | 5.59 | n.s. | 103.56 | 7.51 | n.s. |
| | | 5:1 | 94.29 | 13.13 | n.s. | 98.08 | 11.54 | n.s. | 105.22 | 2.73 | n.s. | 99.11 | 3.52 | n.s. |
| | | 50:1 | 91.76 | 2.19 | n.s. | 95.89 | 16.80 | n.s. | 105.20 | 6.07 | n.s. | 102.99 | 9.18 | n.s. |
| | **3 hrs** | 0 | 100.00 | 0.00 | n.s. | 91.26 | 6.40 | n.s. | 100.00 | 0.00 | n.s. | 84.13 | 8.22 | n.s. |
| | | 0.1:1 | 112.11 | 3.41 | n.s. | 95.09 | 1.97 | n.s. | 105.39 | 7.17 | n.s. | 98.98 | 9.25 | n.s. |
| | | 1:1 | 106.95 | 8.54 | n.s. | 91.58 | 4.10 | n.s. | 108.06 | 8.19 | n.s. | 98.94 | 9.49 | n.s. |
| | | 5:1 | 104.55 | 1.92 | n.s. | 90.77 | 0.13 | n.s. | 107.48 | 10.39 | n.s. | 97.91 | 9.22 | n.s. |
| | | 50:1 | 101.31 | 4.04 | n.s. | 85.59 | 3.20 | n.s. | 103.41 | 7.80 | n.s. | 95.25 | 8.79 | n.s. |
| **CD62L** | **30 min** | 0 | 100.00 | 0.00 | n.s. | 100.90 | 4.06 | n.s. | 100.00 | 0.00 | n.s. | 99.26 | 3.40 | n.s. |
| | | 0.1:1 | 102.66 | 5.90 | n.s. | 103.16 | 5.28 | n.s. | 101.02 | 1.56 | n.s. | 99.68 | 2.27 | n.s. |
| | | 1:1 | 103.06 | 3.58 | n.s. | 102.68 | 5.57 | n.s. | 99.91 | 2.35 | n.s. | 100.00 | 2.31 | n.s. |
| | | 5:1 | 106.80 | 4.07 | n.s. | 100.43 | 2.94 | n.s. | 102.45 | 2.31 | n.s. | 99.30 | 3.31 | n.s. |
| | | 50:1 | 101.46 | 1.85 | n.s. | 99.19 | 5.86 | n.s. | 101.05 | 1.33 | n.s. | 99.98 | 2.87 | n.s. |
| | **3 hrs** | 0 | 100.00 | 0.00 | n.s. | 88.64 | 2.75 | n.s. | 100.00 | 0.00 | n.s. | 98.61 | 3.22 | n.s. |
| | | 0.1:1 | 96.26 | 3.10 | n.s. | 95.52 | 1.61 | n.s. | 98.99 | 4.34 | n.s. | 100.58 | 4.30 | n.s. |
| | | 1:1 | 97.47 | 4.02 | n.s. | 89.93 | 4.27 | n.s. | 100.94 | 4.94 | n.s. | 100.66 | 5.97 | n.s. |
| | | 5:1 | 93.92 | 2.62 | n.s. | 95.65 | 3.81 | n.s. | 101.50 | 3.93 | n.s. | 106.30 | 7.97 | n.s. |
| | | 50:1 | 92.52 | 0.95 | n.s. | 94.43 | 1.34 | n.s. | 101.81 | 3.31 | n.s. | 103.41 | 6.42 | n.s. |

## C)

| | | | S | | | S + αS | | | F | | | F + αF | | |
|---|---|---|---|---|---|---|---|---|---|---|---|---|---|---|
| | | Ratio | % | SEM | *P* | % | SEM | *P* | % | SEM | *P* | % | SEM | *P* |
| CD63 | 30 min | 0 | 100.00 | 0.00 | n.s. | 143.76 | 21.75 | n.s. | 100 | 0 | n.s. | 103.026 | 11.902 | n.s. |
| | | 0.1:1 | 120.80 | 29.23 | n.s. | 118.52 | 21.67 | n.s. | 108.804 | 5.369 | n.s. | 110.3 | 4.8705 | n.s. |
| | | 1:1 | 123.93 | 13.34 | n.s. | 127.88 | 33.37 | n.s. | 108.786 | 12.22 | n.s. | 108.533 | 11.036 | n.s. |
| | | 5:1 | 112.37 | 18.72 | n.s. | 142.27 | 21.02 | n.s. | 96.3122 | 3.0137 | n.s. | 107.647 | 15.166 | n.s. |
| | | 50:1 | 138.49 | 21.47 | n.s. | 148.11 | 28.77 | n.s. | 102.396 | 9.9167 | n.s. | 98.362 | 7.8973 | n.s. |
| | 3 hrs | 0 | 100.00 | 0.00 | n.s. | 112.06 | 13.37 | n.s. | 100 | 0 | n.s. | 107.921 | 9.5234 | n.s. |
| | | 0.1:1 | 126.90 | 22.59 | n.s. | 121.02 | 6.65 | n.s. | 107.717 | 6.2563 | n.s. | 118.489 | 6.8898 | n.s. |
| | | 1:1 | 119.52 | 14.34 | n.s. | 99.83 | 5.84 | n.s. | 110.795 | 8.5524 | n.s. | 108.345 | 8.9374 | n.s. |
| | | 5:1 | 109.17 | 7.86 | n.s. | 125.10 | 8.57 | n.s. | 116.45 | 9.6126 | n.s. | 125.598 | 5.2953 | n.s. |
| | | 50:1 | 123.47 | 20.59 | n.s. | 120.34 | 6.93 | n.s. | 106.755 | 7.0256 | n.s. | 133.025 | 4.6801 | n.s. |
| CD66b | 30 min | 0 | 100.00 | 0.00 | n.s. | 119.91 | 7.15 | n.s. | 100.00 | 0.00 | n.s. | 116.49 | 4.55 | n.s. |
| | | 0.1:1 | 110.41 | 3.11 | n.s. | 106.22 | 6.37 | n.s. | 111.02 | 4.05 | n.s. | 114.14 | 4.08 | n.s. |
| | | 1:1 | 111.64 | 4.34 | n.s. | 111.46 | 11.00 | n.s. | 115.57 | 4.96 | n.s. | 113.16 | 2.54 | n.s. |
| | | 5:1 | 109.96 | 4.28 | n.s. | 110.92 | 3.11 | n.s. | 114.71 | 6.19 | n.s. | 118.36 | 4.55 | n.s. |
| | | 50:1 | 106.24 | 5.65 | n.s. | 116.13 | 8.03 | n.s. | 105.82 | 2.85 | n.s. | 115.66 | 4.69 | n.s. |
| | 3 hrs | 0 | 100.00 | 0.00 | n.s. | 101.54 | 13.03 | n.s. | 100.00 | 0.00 | n.s. | 99.39 | 8.12 | n.s. |
| | | 0.1:1 | 110.63 | 7.27 | n.s. | 112.51 | 1.89 | n.s. | 106.26 | 2.65 | n.s. | 119.46 | 8.58 | n.s. |
| | | 1:1 | 113.19 | 9.24 | n.s. | 95.06 | 15.06 | n.s. | 111.45 | 8.03 | n.s. | 103.89 | 6.29 | n.s. |
| | | 5:1 | 103.43 | 4.32 | n.s. | 106.00 | 5.32 | n.s. | 114.12 | 4.74 | n.s. | 112.96 | 6.00 | n.s. |
| | | 50:1 | 108.40 | 5.03 | n.s. | 109.19 | 3.24 | n.s. | 111.75 | 6.33 | n.s. | 117.04 | 8.36 | n.s. |

## D)

| | | | S | | | S + αS | | | F | | | F + αF | | |
|---|---|---|---|---|---|---|---|---|---|---|---|---|---|---|
| | | Ratio | % | SEM | *P* | % | SEM | *P* | % | SEM | *P* | % | SEM | *P* |
| CD46 | 30 min | 0 | 100.00 | 0.00 | n.s. | 100.13 | 6.93 | n.s. | 100 | 0 | n.s. | 98.3051 | 2.5054 | n.s. |
| | | 0.1:1 | 106.60 | 4.27 | n.s. | 101.95 | 1.87 | n.s. | 99.5759 | 0.7247 | n.s. | 97.4841 | 1.287 | n.s. |
| | | 1:1 | 103.53 | 7.38 | n.s. | 100.69 | 3.29 | n.s. | 97.9448 | 1.4001 | n.s. | 96.9497 | 1.3313 | n.s. |
| | | 5:1 | 99.39 | 4.79 | n.s. | 106.23 | 6.24 | n.s. | 98.2653 | 0.7116 | n.s. | 99.1144 | 1.7814 | n.s. |
| | | 50:1 | 101.29 | 4.51 | n.s. | 103.46 | 4.36 | n.s. | 100.613 | 1.0562 | n.s. | 100.317 | 0.971 | n.s. |
| | 3 hrs | 0 | 100.00 | 0.00 | n.s. | 96.17 | 5.19 | n.s. | 100 | 0 | n.s. | 103.355 | 5.708 | n.s. |
| | | 0.1:1 | 115.26 | 3.86 | n.s. | 115.36 | 6.85 | n.s. | 95.6228 | 3.2042 | n.s. | 104.355 | 2.5145 | n.s. |
| | | 1:1 | 113.65 | 2.84 | n.s. | 101.71 | 3.74 | n.s. | 101.887 | 3.8838 | n.s. | 109.359 | 3.5891 | n.s. |
| | | 5:1 | 107.87 | 5.49 | n.s. | 116.49 | 8.19 | n.s. | 102.167 | 5.5706 | n.s. | 113.114 | 4.7943 | n.s. |
| | | 50:1 | 107.17 | 2.36 | n.s. | 109.95 | 2.57 | n.s. | 106.667 | 3.4991 | n.s. | 108.102 | 4.128 | n.s. |
| CD55 | 30 min | 0 | 100.00 | 0.00 | n.s. | 105.01 | 3.99 | n.s. | 100.00 | 0.00 | n.s. | 103.53 | 3.68 | n.s. |
| | | 0.1:1 | 100.09 | 7.98 | n.s. | 95.95 | 7.80 | n.s. | 101.63 | 1.96 | n.s. | 99.65 | 2.42 | n.s. |
| | | 1:1 | 102.84 | 3.13 | n.s. | 108.79 | 21.26 | n.s. | 101.10 | 2.11 | n.s. | 98.24 | 4.03 | n.s. |
| | | 5:1 | 100.06 | 8.97 | n.s. | 110.77 | 11.56 | n.s. | 101.74 | 3.11 | n.s. | 101.69 | 1.26 | n.s. |
| | | 50:1 | 101.82 | 4.65 | n.s. | 106.16 | 15.94 | n.s. | 101.72 | 4.10 | n.s. | 104.99 | 3.23 | n.s. |
| | 3 hrs | 0 | 100.00 | 0.00 | n.s. | 113.84 | 11.62 | n.s. | 100.00 | 0.00 | n.s. | 98.52 | 5.67 | n.s. |
| | | 0.1:1 | 105.71 | 1.01 | n.s. | 104.19 | 3.54 | n.s. | 96.26 | 0.76 | n.s. | 100.16 | 3.36 | n.s. |
| | | 1:1 | 106.66 | 5.01 | n.s. | 103.16 | 2.24 | n.s. | 98.25 | 3.29 | n.s. | 99.88 | 3.93 | n.s. |
| | | 5:1 | 103.87 | 3.12 | n.s. | 110.53 | 7.01 | n.s. | 102.00 | 4.26 | n.s. | 98.87 | 4.28 | n.s. |
| | | 50:1 | 103.28 | 0.45 | n.s. | 103.21 | 3.61 | n.s. | 99.34 | 5.28 | n.s. | 101.32 | 5.53 | n.s. |
| CD59 | 30 min | 0 | 100.00 | 0.00 | n.s. | 126.65 | 20.25 | n.s. | 100.00 | 0.00 | n.s. | 120.62 | 8.38 | n.s. |
| | | 0.1:1 | 112.70 | 13.51 | n.s. | 110.51 | 14.97 | n.s. | 110.86 | 5.03 | n.s. | 112.62 | 6.14 | n.s. |
| | | 1:1 | 114.58 | 9.31 | n.s. | 145.78 | 50.92 | n.s. | 116.88 | 6.62 | n.s. | 109.32 | 4.07 | n.s. |
| | | 5:1 | 116.56 | 25.99 | n.s. | 136.52 | 33.95 | n.s. | 112.49 | 3.12 | n.s. | 112.11 | 4.99 | n.s. |
| | | 50:1 | 113.64 | 10.40 | n.s. | 135.52 | 40.19 | n.s. | 108.26 | 5.71 | n.s. | 116.81 | 10.18 | n.s. |
| | 3 hrs | 0 | 100.00 | 0.00 | n.s. | 136.32 | 26.70 | n.s. | 100.00 | 0.00 | n.s. | 107.78 | 5.29 | n.s. |
| | | 0.1:1 | 112.25 | 3.24 | n.s. | 105.03 | 2.31 | n.s. | 107.47 | 4.27 | n.s. | 108.15 | 4.61 | n.s. |
| | | 1:1 | 116.10 | 5.45 | n.s. | 104.21 | 3.49 | n.s. | 115.79 | 3.82 | n.s. | 112.63 | 6.45 | n.s. |
| | | 5:1 | 112.50 | 3.02 | n.s. | 114.50 | 12.84 | n.s. | 114.58 | 3.03 | n.s. | 110.71 | 4.71 | n.s. |
| | | 50:1 | 108.83 | 2.17 | n.s. | 107.70 | 2.01 | n.s. | 109.80 | 3.39 | n.s. | 110.70 | 5.52 | n.s. |
| CD93 | 30 min | 0 | 100.00 | 0.00 | n.s. | 119.26 | 2.87 | n.s. | 100.00 | 0.00 | n.s. | 108.26 | 5.27 | n.s. |
| | | 0.1:1 | 127.64 | 12.57 | n.s. | 142.05 | 12.49 | n.s. | 100.57 | 3.08 | n.s. | 102.47 | 6.77 | n.s. |
| | | 1:1 | 119.80 | 8.92 | n.s. | 146.64 | 14.72 | n.s. | 99.95 | 5.17 | n.s. | 99.76 | 4.23 | n.s. |
| | | 5:1 | 147.44 | 8.25 | n.s. | 149.67 | 11.33 | n.s. | 102.46 | 10.70 | n.s. | 105.86 | 10.40 | n.s. |
| | | 50:1 | 121.46 | 9.79 | n.s. | 147.35 | 14.91 | n.s. | 98.14 | 5.58 | n.s. | 103.90 | 5.76 | n.s. |
| | 3 hrs | 0 | 100.00 | 0.00 | n.s. | 124.97 | 5.88 | n.s. | 100.00 | 0.00 | n.s. | 109.63 | 8.48 | n.s. |
| | | 0.1:1 | 116.91 | 5.19 | n.s. | 111.73 | 27.46 | n.s. | 106.54 | 3.95 | n.s. | 105.73 | 4.07 | n.s. |
| | | 1:1 | 110.56 | 8.20 | n.s. | 113.53 | 3.16 | n.s. | 101.05 | 6.37 | n.s. | 111.57 | 7.54 | n.s. |
| | | 5:1 | 114.77 | 28.95 | n.s. | 123.10 | 12.19 | n.s. | 111.36 | 4.30 | n.s. | 108.02 | 3.00 | n.s. |
| | | 50:1 | 96.18 | 6.36 | n.s. | 114.89 | 3.29 | n.s. | 102.52 | 5.27 | n.s. | 110.03 | 6.06 | n.s. |

**Supplementary Table 1**. **Impact of S-nanoparticles and F-nanoparticles alone or pre-coated with antibodies on neutrophil surface marker expression.** Neutrophils were incubated for 30 min or 3 h with the indicated S-nanoparticle-to-cell ratios, either alone or pre-coated with a monoclonal anti-S antibody (αS; **Table 1**). F-nanoparticles, with or without pre-coating with anti-F antibody (αF; **Table 1**), were used as comparators. Surface markers were stained with fluorophore-conjugated antibodies and analyzed by flow cytometry (**Table 2**). Markers were selected for the monitoring of the following parameters: (**A)** Interactions with immune complexes (CD16, CD32, CD64); (**B)** Adhesion (CD11b, CD15, CD62L); (**C)** Degranulation of primary (CD63) and secondary (CD66b) granules; and (**D)** Complement regulation (CD46, CD55, CD59, CD93). Results were expressed as percent change in the mean fluorescence intensity (MFI) relative to non-stimulated cells. Data are shown as mean ± SEM (n = 3 donors). n.s.: non-significant.
