## Supplemental Table 2 for "No Evidence of Direct Activation of Human Neutrophil Responses by Multivalent Prefusion Trimeric SARS-CoV-2 Spike Protein *ex vivo*"

## A)

| | | | SCoV2 | | | SCoV2 + αS | | | | RSV | | | RSV + αF | | |
|---|---|---|---|---|---|---|---|---|---|---|---|---|---|---|---|
| | | Ratio | % | SEM | *P* | % | SEM | *P* | Ratio | % | SEM | *P* | % | SEM | *P* |
| CD16 | 30 min | 0 | 100,00 | 0,00 | n.s. | 96,83 | 2,34 | n.s. | 0 | 100,00 | 0,00 | n.s. | 99,66 | 1,03 | n.s. |
| | | 0.001:1 | 110,09 | 2,62 | n.s. | 100,68 | 2,44 | n.s. | 0.000025:1 | 102,10 | 1,19 | n.s. | 96,28 | 1,46 | n.s. |
| | | 0.1:1 | 104,93 | 7,04 | n.s. | 100,43 | 3,98 | n.s. | 0.0025:1 | 109,64 | 3,99 | n.s. | 99,46 | 2,64 | n.s. |
| | | 1:1 | 105,69 | 7,05 | n.s. | 102,32 | 4,31 | n.s. | 0.025:1 | 108,78 | 4,73 | n.s. | 105,50 | 2,01 | n.s. |
| | | 10:1 | 101,82 | 3,80 | n.s. | 99,40 | 3,82 | n.s. | 0.25:1 | 111,12 | 7,20 | n.s. | 109,73 | 2,10 | n.s. |
| | 3 hrs | 0 | 100,00 | 0,00 | n.s. | 96,64 | 6,49 | n.s. | 0 | 100,00 | 0,00 | n.s. | 105,47 | 5,57 | n.s. |
| | | 0.001:1 | 101,05 | 12,55 | n.s. | 79,54 | 3,18 | n.s. | 0.000025:1 | 101,75 | 2,19 | n.s. | 91,87 | 8,57 | n.s. |
| | | 0.1:1 | 101,01 | 9,19 | n.s. | 98,54 | 9,78 | n.s. | 0.0025:1 | 101,67 | 6,31 | n.s. | 102,88 | 5,18 | n.s. |
| | | 1:1 | 99,84 | 7,28 | n.s. | 99,58 | 9,23 | n.s. | 0.025:1 | 84,97 | 17,60 | n.s. | 100,61 | 6,90 | n.s. |
| | | 10:1 | 98,90 | 5,56 | n.s. | 95,93 | 7,22 | n.s. | 0.25:1 | 88,66 | 22,85 | n.s. | 104,63 | 8,34 | n.s. |
| CD32 | 30 min | 0 | 100,00 | 0,00 | n.s. | 96,65 | 2,39 | n.s. | 0 | 100,00 | 0,00 | n.s. | 107,71 | 3,60 | n.s. |
| | | 0.001:1 | 111,78 | 4,92 | n.s. | 104,11 | 3,77 | n.s. | 0.000025:1 | 101,43 | 1,71 | n.s. | 97,42 | 2,87 | n.s. |
| | | 0.1:1 | 111,09 | 3,26 | n.s. | 102,55 | 1,89 | n.s. | 0.0025:1 | 109,29 | 5,50 | n.s. | 98,10 | 2,51 | n.s. |
| | | 1:1 | 111,33 | 0,70 | n.s. | 105,75 | 3,34 | n.s. | 0.025:1 | 105,34 | 0,97 | n.s. | 109,42 | 6,24 | n.s. |
| | | 10:1 | 108,40 | 4,14 | n.s. | 105,79 | 5,35 | n.s. | 0.25:1 | 104,63 | 1,66 | n.s. | 101,42 | 0,33 | n.s. |
| | 3 hrs | **0** | **100,00** | **0,00** | **n.s.** | **130,59** | **5,92** | **n.s.** | 0 | 100,00 | 0,00 | n.s. | 104,86 | 4,94 | n.s. |
| | | **0.001:1** | **132,48** | **8,11** | **n.s.** | **111,83** | **24,27** | **n.s.** | 0.000025:1 | 106,42 | 4,81 | n.s. | 93,14 | 4,73 | n.s. |
| | | **0.1:1** | **155,82** | **15,75** | **0,0060** | **143,17** | **11,66** | **n.s.** | 0.0025:1 | 112,14 | 8,67 | n.s. | 107,88 | 4,71 | n.s. |
| | | **1:1** | **149,09** | **16,81** | **n.s.** | **145,85** | **9,12** | **n.s.** | 0.025:1 | 92,29 | 9,32 | n.s. | 100,49 | 3,64 | n.s. |
| | | **10:1** | **150,77** | **12,43** | **n.s.** | **150,03** | **15,73** | **n.s.** | 0.25:1 | 76,63 | 20,50 | n.s. | 100,22 | 5,88 | n.s. |
| CD64 | 30 min | 0 | 100,00 | 0,00 | n.s. | 96,97 | 0,94 | n.s. | 0 | 100,00 | 0,00 | n.s. | 99,91 | 0,55 | n.s. |
| | | 0.001:1 | 94,00 | 7,96 | n.s. | 93,45 | 6,63 | n.s. | 0.000025:1 | 92,99 | 2,70 | n.s. | 94,31 | 0,93 | n.s. |
| | | 0.1:1 | 90,13 | 5,57 | n.s. | 94,02 | 3,94 | n.s. | 0.0025:1 | 99,77 | 6,51 | n.s. | 99,10 | 6,72 | n.s. |
| | | 1:1 | 92,33 | 4,35 | n.s. | 94,85 | 8,62 | n.s. | 0.025:1 | 87,53 | 2,28 | n.s. | 109,52 | 12,65 | n.s. |
| | | 10:1 | 88,63 | 1,11 | n.s. | 91,01 | 4,41 | n.s. | 0.25:1 | 89,60 | 1,18 | n.s. | 91,05 | 1,39 | n.s. |
| | 3 hrs | 0 | 100,00 | 0,00 | n.s. | 102,50 | 11,32 | n.s. | 0 | 100,00 | 0,00 | n.s. | 100,13 | 3,99 | n.s. |
| | | 0.001:1 | 102,77 | 21,00 | n.s. | 91,46 | 0,64 | n.s. | 0.000025:1 | 114,40 | 0,89 | n.s. | 105,17 | 1,34 | n.s. |
| | | 0.1:1 | 103,03 | 10,69 | n.s. | 95,82 | 10,63 | n.s. | 0.0025:1 | 117,42 | 8,70 | n.s. | 102,38 | 3,36 | n.s. |
| | | 1:1 | 98,84 | 9,00 | n.s. | 98,93 | 15,10 | n.s. | 0.025:1 | 103,11 | 8,34 | n.s. | 107,79 | 2,68 | n.s. |
| | | 10:1 | 108,41 | 11,24 | n.s. | 98,02 | 9,13 | n.s. | 0.25:1 | 78,55 | 8,95 | n.s. | 108,47 | 5,26 | n.s. |

## B)

| | | | SCoV2 | | | SCoV2 + αS | | | | RSV | | | RSV + αF | | |
|---|---|---|---|---|---|---|---|---|---|---|---|---|---|---|---|
| | | Ratio | % | SEM | *P* | % | SEM | *P* | Ratio | % | SEM | *P* | % | SEM | *P* |
| CD11b | 30 min | 0 | 100.00 | 0.00 | n.s. | 107.01 | 8.18 | n.s. | 0 | 100 | 0 | n.s. | 108.427 | 4.2149 | n.s. |
| | | 0.001:1 | 104.85 | 7.71 | n.s. | 100.35 | 6.88 | n.s. | 0.000025:1 | 94.8155 | 7.5735 | n.s. | 93.8424 | 4.44826 | n.s. |
| | | 0.1:1 | 105.48 | 2.98 | n.s. | 101.67 | 4.51 | n.s. | 0.0025:1 | 106.644 | 7.0066 | n.s. | 92.4541 | 4.12396 | n.s. |
| | | 1:1 | 106.07 | 4.98 | n.s. | 102.12 | 5.53 | n.s. | 0.025:1 | 103.635 | 3.476 | n.s. | 103.076 | 4.10819 | n.s. |
| | | 10:1 | 110.86 | 2.41 | n.s. | 112.38 | 3.43 | n.s. | 0.25:1 | 109.408 | 1.9596 | n.s. | 103.491 | 2.70218 | n.s. |
| | 3 hrs | **0** | **100.00** | **0.00** | **n.s.** | **131.37** | **5.92** | **n.s.** | 0 | 100 | 0 | n.s. | 118.224 | 7.38098 | n.s. |
| | | **0.001:1** | **133.10** | **0.00** | **n.s.** | **107.07** | **24.12** | **n.s.** | 0.000025:1 | 111.188 | 9.5656 | n.s. | 106.918 | 13.2619 | n.s. |
| | | **0.1:1** | **144.12** | **0.04** | **0.0322** | **144.81** | **4.47** | **n.s.** | 0.0025:1 | 122.213 | 14.759 | n.s. | 110.44 | 7.56242 | n.s. |
| | | **1:1** | **133.06** | **6.18** | **n.s.** | **133.00** | **0.81** | **n.s.** | 0.025:1 | 101.648 | 17.209 | n.s. | 108.773 | 8.79501 | n.s. |
| | | **10:1** | **153.64** | **0.12** | **0.0060** | **145.72** | **5.58** | **n.s.** | 0.25:1 | 93.4773 | 25.303 | n.s. | 111.55 | 12.2775 | n.s. |
| CD15 | 30 min | 0 | 100.00 | 0.00 | n.s. | 94.60 | 0.97 | n.s. | 0 | 100.00 | 0.00 | n.s. | 107.61 | 7.00 | n.s. |
| | | 0.001:1 | 92.52 | 1.01 | n.s. | 94.49 | 1.28 | n.s. | 0.000025:1 | 95.90 | 5.45 | n.s. | 100.09 | 1.90 | n.s. |
| | | 0.1:1 | 91.45 | 6.72 | n.s. | 97.15 | 1.81 | n.s. | 0.0025:1 | 91.12 | 3.36 | n.s. | 96.53 | 4.22 | n.s. |
| | | 1:1 | 86.93 | 5.49 | n.s. | 95.47 | 2.45 | n.s. | 0.025:1 | 87.43 | 6.43 | n.s. | 100.77 | 6.20 | n.s. |
| | | 10:1 | 91.32 | 2.42 | n.s. | 91.32 | 4.48 | n.s. | 0.25:1 | 88.07 | 6.52 | n.s. | 100.74 | 7.87 | n.s. |
| | 3 hrs | 0 | 100.00 | 0.00 | n.s. | 92.04 | 9.93 | n.s. | 0 | 100.00 | 0.00 | n.s. | 97.69 | 5.11 | n.s. |
| | | 0.001:1 | 101.28 | 5.71 | n.s. | 85.96 | 17.10 | n.s. | 0.000025:1 | 93.76 | 3.82 | n.s. | 93.04 | 4.93 | n.s. |
| | | 0.1:1 | 93.50 | 1.78 | n.s. | 105.95 | 3.58 | n.s. | 0.0025:1 | 91.04 | 4.10 | n.s. | 98.92 | 1.32 | n.s. |
| | | 1:1 | 109.27 | 9.84 | n.s. | 98.95 | 7.49 | n.s. | 0.025:1 | 82.84 | 1.82 | n.s. | 93.34 | 5.90 | n.s. |
| | | 10:1 | 94.70 | 18.60 | n.s. | 94.20 | 10.22 | n.s. | 0.25:1 | 84.73 | 0.43 | n.s. | 86.16 | 8.97 | n.s. |
| CD62L | 30 min | **0** | **100.00** | **0.00** | **n.s.** | **102.12** | **0.95** | **n.s.** | 0 | 100.00 | 0.00 | n.s. | 100.07 | 1.41 | n.s. |
| | | **0.001:1** | **109.36** | **1.26** | **n.s.** | **104.73** | **6.07** | **n.s.** | 0.000025:1 | 106.43 | 2.81 | n.s. | 102.83 | 1.50 | n.s. |
| | | **0.1:1** | **106.77** | **2.94** | **n.s.** | **128.42** | **26.20** | **n.s.** | 0.0025:1 | 109.25 | 2.67 | n.s. | 101.87 | 1.73 | n.s. |
| | | **1:1** | **109.80** | **1.93** | **0.0322** | **136.66** | **30.44** | **n.s.** | 0.025:1 | 109.33 | 2.74 | n.s. | 110.42 | 2.18 | n.s. |
| | | **10:1** | **112.04** | **1.86** | **0.0060** | **145.05** | **29.67** | **n.s.** | 0.25:1 | 116.59 | 3.53 | n.s. | 107.53 | 7.87 | n.s. |
| | 3 hrs | 0 | 100.00 | 0.00 | n.s. | 126.98 | 24.84 | n.s. | **0** | **100.00** | **0.00** | **n.s.** | **105.73** | **3.30** | **n.s.** |
| | | 0.001:1 | 105.74 | 1.34 | n.s. | 102.72 | 1.27 | n.s. | **0.000025:1** | **104.46** | **1.95** | **n.s.** | **96.07** | **6.30** | **n.s.** |
| | | 0.1:1 | 104.79 | 2.51 | n.s. | 131.99 | 30.04 | n.s. | **0.0025:1** | **105.22** | **3.11** | **n.s.** | **98.88** | **3.45** | **n.s.** |
| | | 1:1 | 104.60 | 1.56 | n.s. | 136.77 | 34.25 | n.s. | **0.025:1** | **104.94** | **4.08** | **n.s.** | **111.21** | **3.59** | **n.s.** |
| | | 10:1 | 107.68 | 2.69 | n.s. | 139.20 | 29.27 | n.s. | **0.25:1** | **126.59** | **1.74** | **0.0427** | **127.70** | **4.81** | **n.s.** |

**C)**

| | | | SCoV2 | | | SCoV2 + αS | | | | RSV | | | RSV + αF | | |
|---|---|---|---|---|---|---|---|---|---|---|---|---|---|---|---|
| | | Ratio | % | SEM | *P* | % | SEM | *P* | Ratio | % | SEM | *P* | % | SEM | *P* |
| **CD63** | **30 min** | 0 | 100.00 | 0.00 | n.s. | 87.69 | 4.92 | n.s. | 0 | 100 | 0 | n.s. | 93.6645 | 19.5643 | n.s. |
| | | 0.001:1 | 101.19 | 5.99 | n.s. | 114.84 | 10.45 | n.s. | 0.000025:1 | 83.7628 | 18.241 | n.s. | 85.0185 | 14.6454 | n.s. |
| | | 0.1:1 | 90.22 | 13.90 | n.s. | 90.84 | 7.36 | n.s. | 0.0025:1 | 87.5847 | 25.549 | n.s. | 93.9769 | 19.989 | n.s. |
| | | 1:1 | 88.88 | 11.70 | n.s. | 91.23 | 12.61 | n.s. | 0.025:1 | 86.4297 | 18.696 | n.s. | 92.0394 | 21.4898 | n.s. |
| | | 10:1 | 93.13 | 11.01 | n.s. | 99.68 | 14.60 | n.s. | 0.25:1 | 90.6319 | 6.5055 | n.s. | 83.0856 | 27.9241 | n.s. |
| | **3 hrs** | 0 | 100.00 | 0.00 | n.s. | 114.20 | 9.63 | n.s. | 0 | 100 | 0 | n.s. | 106.446 | 5.11134 | n.s. |
| | | 0.001:1 | 104.56 | 1.26 | n.s. | 100.94 | 1.17 | n.s. | 0.000025:1 | 106.622 | 6.3973 | n.s. | 99.1593 | 4.46916 | n.s. |
| | | 0.1:1 | 114.63 | 21.60 | n.s. | 114.19 | 10.28 | n.s. | 0.0025:1 | 97.6139 | 7.5137 | n.s. | 95.1866 | 5.60509 | n.s. |
| | | 1:1 | 113.61 | 13.32 | n.s. | 106.48 | 8.89 | n.s. | 0.025:1 | 107.219 | 10.946 | n.s. | 105.006 | 8.53763 | n.s. |
| | | 10:1 | 134.89 | 26.79 | n.s. | 116.65 | 14.50 | n.s. | 0.25:1 | 91.5985 | 4.3958 | n.s. | 99.9132 | 3.78749 | n.s. |
| **CD66b** | **30 min** | 0 | 100.00 | 0.00 | n.s. | 96.51 | 2.03 | n.s. | 0 | 100.00 | 0.00 | n.s. | 107.58 | 6.03 | n.s. |
| | | 0.001:1 | 105.50 | 7.45 | n.s. | 103.71 | 2.95 | n.s. | 0.000025:1 | 103.74 | 3.03 | n.s. | 105.63 | 2.15 | n.s. |
| | | 0.1:1 | 98.35 | 4.77 | n.s. | 99.99 | 2.28 | n.s. | 0.0025:1 | 105.72 | 2.98 | n.s. | 106.30 | 1.61 | n.s. |
| | | 1:1 | 101.80 | 6.19 | n.s. | 100.62 | 7.49 | n.s. | 0.025:1 | 111.27 | 3.27 | n.s. | 105.93 | 3.29 | n.s. |
| | | 10:1 | 99.19 | 5.28 | n.s. | 98.79 | 3.56 | n.s. | 0.25:1 | 97.45 | 5.99 | n.s. | 87.32 | 19.25 | n.s. |
| | **3 hrs** | 0 | 100.00 | 0.00 | n.s. | 107.17 | 4.56 | n.s. | 0 | 100.00 | 0.00 | n.s. | 103.98 | 1.50 | n.s. |
| | | 0.001:1 | 108.98 | 2.32 | n.s. | 108.17 | 7.45 | n.s. | 0.000025:1 | 102.65 | 1.94 | n.s. | 104.50 | 2.42 | n.s. |
| | | 0.1:1 | 116.68 | 4.62 | n.s. | 109.36 | 2.52 | n.s. | 0.0025:1 | 97.69 | 13.35 | n.s. | 102.01 | 2.09 | n.s. |
| | | 1:1 | 123.87 | 8.87 | n.s. | 106.62 | 3.99 | n.s. | 0.025:1 | 110.86 | 4.53 | n.s. | 106.20 | 1.10 | n.s. |
| | | 10:1 | 124.55 | 3.32 | n.s. | 109.74 | 3.40 | n.s. | 0.25:1 | 103.17 | 2.84 | n.s. | 103.15 | 2.51 | n.s. |

**D)**

| | | | SCoV2 | | | SCoV2 + αS | | | | RSV | | | RSV + αF | | |
|---|---|---|---|---|---|---|---|---|---|---|---|---|---|---|---|
| | | Ratio | % | SEM | *P* | % | SEM | *P* | Ratio | % | SEM | *P* | % | SEM | *P* |
| **CD46** | **30 min** | 0 | 100.00 | 0.00 | n.s. | 101.09 | 3.72 | n.s. | 0 | 100 | 0 | n.s. | 103.687 | 1.93079 | n.s. |
| | | 0.001:1 | 114.20 | 7.24 | n.s. | 110.75 | 6.37 | n.s. | 0.000025:1 | 103.445 | 0.3231 | n.s. | 101.274 | 0.6481 | n.s. |
| | | 0.1:1 | 113.76 | 8.94 | n.s. | 106.59 | 5.83 | n.s. | 0.0025:1 | 104.311 | 0.3212 | n.s. | 100.143 | 0.88168 | n.s. |
| | | 1:1 | 107.69 | 5.54 | n.s. | 105.41 | 7.62 | n.s. | 0.025:1 | 104.778 | 0.7697 | n.s. | 106.682 | 3.79829 | n.s. |
| | | 10:1 | 105.29 | 5.86 | n.s. | 101.20 | 7.42 | n.s. | 0.25:1 | 110.35 | 1.3682 | **0.0075** | 108.388 | 0.79776 | **0.0356** |
| | **3 hrs** | 0 | 100.00 | 0.00 | n.s. | 89.73 | 16.43 | n.s. | 0 | 100 | 0 | n.s. | 106.132 | 1.9177 | n.s. |
| | | 0.001:1 | 109.70 | 1.05 | n.s. | 92.09 | 8.58 | n.s. | 0.000025:1 | 103.522 | 2.5156 | n.s. | 94.812 | 3.0907 | n.s. |
| | | 0.1:1 | 87.59 | 18.33 | n.s. | 92.10 | 15.03 | n.s. | 0.0025:1 | 104.677 | 4.0954 | n.s. | 96.849 | 3.86379 | n.s. |
| | | 1:1 | 96.91 | 16.92 | n.s. | 95.05 | 11.70 | n.s. | 0.025:1 | 97.8392 | 0.4832 | n.s. | 105.702 | 5.2867 | n.s. |
| | | 10:1 | 97.02 | 11.97 | n.s. | 89.83 | 12.52 | n.s. | 0.25:1 | 124.724 | 6.0929 | n.s. | 124.202 | 7.23374 | n.s. |
| **CD55** | **30 min** | 0 | 100.00 | 0.00 | n.s. | 95.12 | 3.35 | n.s. | 0 | 100.00 | 0.00 | n.s. | 101.78 | 2.82 | n.s. |
| | | 0.001:1 | 102.66 | 3.13 | n.s. | 103.51 | 3.04 | n.s. | 0.000025:1 | 95.67 | 5.24 | n.s. | 97.00 | 3.79 | n.s. |
| | | 0.1:1 | 101.44 | 6.65 | n.s. | 97.40 | 5.32 | n.s. | 0.0025:1 | 94.67 | 3.71 | n.s. | 92.00 | 4.93 | n.s. |
| | | 1:1 | 95.85 | 5.64 | n.s. | 94.51 | 7.63 | n.s. | 0.025:1 | 95.33 | 3.48 | n.s. | 96.00 | 7.02 | n.s. |
| | | 10:1 | 93.92 | 7.38 | n.s. | 95.48 | 6.52 | n.s. | 0.25:1 | 98.67 | 1.76 | n.s. | 95.33 | 3.53 | n.s. |
| | **3 hrs** | 0 | 100.00 | 0.00 | n.s. | 97.46 | 4.39 | n.s. | 0 | 100.00 | 0.00 | n.s. | 100.51 | 8.45 | n.s. |
| | | 0.001:1 | 106.34 | 5.19 | n.s. | 79.71 | 13.75 | n.s. | 0.000025:1 | 93.33 | 6.36 | n.s. | 91.67 | 8.57 | n.s. |
| | | 0.1:1 | 98.82 | 5.51 | n.s. | 101.05 | 2.94 | n.s. | 0.0025:1 | 95.33 | 8.67 | n.s. | 93.33 | 4.67 | n.s. |
| | | 1:1 | 105.14 | 6.13 | n.s. | 100.45 | 3.56 | n.s. | 0.025:1 | 90.33 | 8.25 | n.s. | 91.67 | 7.84 | n.s. |
| | | 10:1 | 103.40 | 3.21 | n.s. | 95.58 | 5.31 | n.s. | 0.25:1 | 96.67 | 9.02 | n.s. | 96.00 | 11.24 | n.s. |
| **CD59** | **30 min** | 0 | 100.00 | 0.00 | n.s. | 109.76 | 0.91 | n.s. | 0 | 100.00 | 0.00 | n.s. | 120.29 | 12.45 | n.s. |
| | | 0.001:1 | 91.34 | 13.21 | n.s. | 97.64 | 2.55 | n.s. | 0.000025:1 | 101.46 | 6.21 | n.s. | 103.80 | 3.20 | n.s. |
| | | 0.1:1 | 101.76 | 4.70 | n.s. | 106.10 | 3.66 | n.s. | 0.0025:1 | 95.13 | 4.41 | n.s. | 100.45 | 2.38 | n.s. |
| | | 1:1 | 91.67 | 9.28 | n.s. | 97.82 | 6.43 | n.s. | 0.025:1 | 90.38 | 4.47 | n.s. | 110.79 | 12.04 | n.s. |
| | | 10:1 | 106.72 | 3.74 | n.s. | 95.05 | 4.66 | n.s. | 0.25:1 | 97.45 | 5.54 | n.s. | 96.42 | 2.01 | n.s. |
| | **3 hrs** | 0 | 100.00 | 0.00 | n.s. | 108.53 | 5.97 | n.s. | 0 | 100.00 | 0.00 | n.s. | 91.75 | 6.74 | n.s. |
| | | 0.001:1 | 111.52 | 2.81 | n.s. | 91.21 | 6.35 | n.s. | 0.000025:1 | 87.92 | 15.69 | n.s. | 95.37 | 16.95 | n.s. |
| | | 0.1:1 | 105.86 | 11.93 | n.s. | 117.81 | 12.91 | n.s. | 0.0025:1 | 85.94 | 16.08 | n.s. | 102.90 | 18.94 | n.s. |
| | | 1:1 | 127.60 | 2.39 | n.s. | 121.83 | 8.63 | n.s. | 0.025:1 | 81.33 | 11.80 | n.s. | 89.46 | 18.84 | n.s. |
| | | 10:1 | 111.05 | 20.23 | n.s. | 123.44 | 18.34 | n.s. | 0.25:1 | 93.26 | 24.13 | n.s. | 84.88 | 16.16 | n.s. |
| **CD93** | **30 min** | 0 | 100.00 | 0.00 | n.s. | 100.96 | 10.19 | n.s. | 0 | 100.00 | 0.00 | n.s. | 98.38 | 7.69 | n.s. |
| | | 0.001:1 | 96.80 | 0.82 | n.s. | 122.99 | 11.90 | n.s. | 0.000025:1 | 97.25 | 7.51 | n.s. | 106.60 | 12.87 | n.s. |
| | | 0.1:1 | 100.67 | 7.88 | n.s. | 123.38 | 2.95 | n.s. | 0.0025:1 | 106.44 | 11.99 | n.s. | 97.14 | 7.32 | n.s. |
| | | 1:1 | 95.23 | 4.87 | n.s. | 113.06 | 10.31 | n.s. | 0.025:1 | 90.34 | 9.81 | n.s. | 116.03 | 20.14 | n.s. |
| | | 10:1 | 92.11 | 13.08 | n.s. | 143.66 | 6.70 | n.s. | 0.25:1 | 101.35 | 2.94 | n.s. | 107.84 | 3.46 | n.s. |
| | **3 hrs** | 0 | 100.00 | 0.00 | n.s. | 117.00 | 6.71 | n.s. | 0 | 100.00 | 0.00 | n.s. | 107.03 | 4.55 | n.s. |
| | | 0.001:1 | 82.41 | 2.38 | n.s. | 122.87 | 30.92 | n.s. | 0.000025:1 | 122.19 | 12.19 | n.s. | 112.02 | 12.85 | n.s. |
| | | 0.1:1 | 97.57 | 7.22 | n.s. | 114.58 | 4.40 | n.s. | 0.0025:1 | 138.41 | 15.12 | n.s. | 112.08 | 4.89 | n.s. |
| | | 1:1 | 99.10 | 7.08 | n.s. | 119.07 | 9.45 | n.s. | 0.025:1 | 119.33 | 16.51 | n.s. | 108.15 | 21.00 | n.s. |
| | | 10:1 | 87.70 | 1.77 | n.s. | 119.36 | 9.88 | n.s. | 0.25:1 | 87.90 | 13.01 | n.s. | 126.49 | 11.77 | n.s. |

**Supplementary Table 2**. **Impact of Inactivated-SARS-CoV-2 and -RSV alone or pre-coated with antibodies on neutrophil surface marker expression.** Neutrophils were incubated for 30 min or 3 h with the indicated BPL-inactivated SARS-CoV-2 (SCoV2)-to-cell ratios, either alone or pre-coated with a monoclonal anti-S antibody (αS; **Table 1**). BPL-inactivated RSV, with or without pre-coating with anti-F antibody (αF; **Table 1**), were used as comparators. Surface markers were stained with fluorophore-conjugated antibodies and analyzed by flow cytometry (**Table 2**). Markers were selected for the monitoring of the following parameters: (**A)** Interactions with immune complexes (CD16, CD32, CD64); (**B)** Adhesion (CD11b, CD15, CD62L); (**C)** Degranulation of primary (CD63) and secondary (CD66b) granules; and (**D)** Complement regulation (CD46, CD55, CD59, CD93). Results were expressed as percent change in the mean fluorescence intensity (MFI) relative to non-stimulated cells. Data are shown as mean ± SEM (n = 3 donors). When statistically different from non-stimulated cells, *P* values are indicated in Bold. n.s.: non-significant.
