## Supplemental Table 3 for "No Evidence of Direct Activation of Human Neutrophil Responses by Multivalent Prefusion Trimeric SARS-CoV-2 Spike Protein *ex vivo*"

| | | S | | | | | SCoV2 | | | | |
|---|---|---|---|---|---|---|---|---|---|---|---|
| | | + αS | | + αS Mix | | | + αS | | + αS Mix | | |
| | | % | SEM | % | SEM | *P* | % | SEM | % | SEM | *P* |
| **CD16** | 30 min | 109.23 | 3.71 | 102.13 | 0.48 | n.s. | 95.84 | 9.11 | 91.89 | 8.00 | n.s. |
| | 3 hrs | 107.09 | 1.86 | 100.92 | 3.54 | n.s. | 98.94 | 8.96 | 97.16 | 6.97 | n.s. |
| **CD32** | 30 min | 102.09 | 3.26 | 99.89 | 2.59 | n.s. | 106.21 | 7.38 | 110.18 | 11.62 | n.s. |
| | 3 hrs | 101.83 | 1.43 | 95.40 | 1.82 | n.s. | 106.34 | 5.05 | 95.65 | 5.16 | n.s. |
| **CD64** | 30 min | 96.03 | 0.00 | 80.92 | 0.00 | – | 87.63 | 0.00 | 75.30 | 0.00 | – |
| | 3 hrs | 99.82 | 0.00 | 98.63 | 0.00 | – | 99.03 | 0.00 | 101.83 | 0.00 | – |
| **CD11b** | 30 min | 101.84 | 8.89 | 102.83 | 4.32 | n.s. | 125.68 | 13.12 | 135.50 | 12.20 | n.s. |
| | 3 hrs | 99.96 | 7.30 | 96.69 | 2.94 | n.s. | 135.86 | 7.99 | 121.91 | 9.06 | n.s. |
| **CD15** | 30 min | 92.68 | 5.38 | 92.14 | 8.07 | n.s. | 89.84 | 4.82 | 94.97 | 9.64 | n.s. |
| | 3 hrs | 93.45 | 2.16 | 86.88 | 6.96 | n.s. | 90.48 | 2.58 | 84.46 | 6.69 | n.s. |
| **CD62L** | 30 min | 103.48 | 0.54 | 103.67 | 1.08 | n.s. | 91.50 | 12.99 | 90.65 | 10.97 | n.s. |
| | 3 hrs | 101.79 | 5.50 | 94.56 | 4.81 | n.s. | 113.27 | 1.51 | 113.84 | 2.00 | n.s. |
| **CD63** | 30 min | 91.11 | 9.09 | 105.58 | 11.19 | n.s. | 93.53 | 1.66 | 89.69 | 11.25 | n.s. |
| | 3 hrs | 97.04 | 4.95 | 88.23 | 8.24 | n.s. | 116.35 | 16.59 | 95.69 | 17.95 | n.s. |
| **CD66b** | 30 min | 98.16 | 3.52 | 100.07 | 3.02 | n.s. | 103.07 | 1.57 | 111.10 | 2.34 | n.s. |
| | 3 hrs | 107.69 | 4.56 | 93.92 | 1.74 | n.s. | 109.06 | 4.50 | 93.79 | 4.34 | n.s. |
| **CD46** | 30 min | 104.56 | 0.41 | 100.25 | 1.39 | n.s. | 88.44 | 15.16 | 85.08 | 11.04 | n.s. |
| | 3 hrs | 104.87 | 5.10 | 99.05 | 2.50 | n.s. | 104.90 | 12.50 | 98.66 | 2.96 | n.s. |
| **CD55** | 30 min | 99.62 | 0.26 | 97.40 | 3.32 | n.s. | 96.80 | 3.00 | 96.03 | 2.78 | n.s. |
| | 3 hrs | 105.87 | 4.43 | 96.45 | 2.76 | n.s. | 97.93 | 8.83 | 94.78 | 4.08 | n.s. |
| **CD59** | 30 min | 100.07 | 0.33 | 92.75 | 7.26 | n.s. | 108.35 | 1.15 | 116.78 | 20.73 | n.s. |
| | 3 hrs | 90.93 | 0.14 | 88.56 | 7.63 | n.s. | 96.35 | 4.64 | 93.33 | 5.49 | n.s. |
| **CD93** | 30 min | 115.09 | 2.20 | 90.21 | 3.66 | n.s. | 101.75 | 9.87 | 85.98 | 8.84 | n.s. |
| | 3 hrs | 104.21 | 10.57 | 93.16 | 8.45 | n.s. | 95.98 | 6.20 | 92.59 | 9.16 | n.s. |

**Supplementary Table 3**. **Impact of pre-coating of nanoparticles or inactivated virus with a single antibody vs a mix of five antibodies on neutrophil surface marker expression.** Neutrophils were incubated for 30 min or 3 h with S-nanoparticles (S; ratio S-nanoparticle : neutrophil = 50:1) or BPL-inactivated SARS-CoV-2 (SCoV2; ration SCoV2 : neutrophil = 10:1) either alone or pre-coated with single a monoclonal anti-S antibody (αS; **Table 1**) or a mixture of five monoclonal antibodies (αSmix; **Table 1**). The indicated surface markers were stained with fluorophore-conjugated antibodies and analyzed by flow cytometry (**Table 2**). Results were expressed as percent change in the mean fluorescence intensity (MFI) relative to cells incubated with antibodies alone. Data are shown as mean ± SEM (n = 3 donors). n.s.: non-significant.
